# Prioritizing genomic variants through neuro-symbolic, knowledge-enhanced learning

**DOI:** 10.1101/2023.11.08.566179

**Authors:** Azza Althagafi, Fernando Zhapa-Camacho, Robert Hoehndorf

## Abstract

**Motivation:** Whole-exome and genome sequencing have become common tools in diagnosing patients with rare diseases. Despite their success, this approach leaves many patients undiagnosed. A common argument is that more disease variants still await discovery, or the novelty of disease phenotypes results from a combination of variants in multiple disease-related genes. Interpreting the phenotypic consequences of genomic variants relies on information about gene functions, gene expression, physiology, and other genomic features. Phenotype-based methods to identify variants involved in genetic diseases combine molecular features with prior knowledge about the phenotypic consequences of altering gene functions. While phenotype-based methods have been successfully applied to prioritizing variants, such methods are based on known gene–disease or gene–phenotype associations as training data and are applicable to genes that have phenotypes associated, thereby limiting their scope. In addition, phenotypes are not assigned uniformly by different clinicians, and phenotype-based methods need to account for this variability.

**Results:** We developed an Embedding-based Phenotype Variant Predictor (EmbedPVP), a computational method to prioritize variants involved in genetic diseases by combining genomic information and clinical phenotypes. EmbedPVP leverages a large amount of background knowledge from human and model organisms about molecular mechanisms through which abnormal phenotypes may arise. Specifically, EmbedPVP incorporates phenotypes linked to genes, functions of gene products, and the anatomical site of gene expression, and systematically relates them to their phenotypic effects through neuro-symbolic, knowledge-enhanced machine learning. We demonstrate EmbedPVP’s efficacy on a large set of synthetic genomes and genomes matched with clinical information.

**Availability:** EmbedPVP and all evaluation experiments are freely available at https://github.com/bio-ontology-research-group/EmbedPVP.

**Supplementary information:** Supplementary data are available at *Bioinformatics*.

## 1 Introduction

The contribution of genetics to human diseases ranges from almost 100% for monogenic, Mendelian disorders to much smaller percentages for complex diseases, including infectious disease Hyman (2000). Understanding how variation in an individual’s genome relates to disease risk is important, as it allows us to prevent and predict negative health effects in individuals, generate better diagnoses and prognoses for disease, and enable new approaches for treatment and development of new drugs Bloss *et al*. (2011). Predicting possible health effects from genome sequences is a significant emerging challenge and is important to support genetic counseling and prevent major health problems. Whole Exome and Genome Sequencing (WGS/WES) has become a common tool in the diagnosis of patients with rare diseases as it has improved diagnostic yields and enables efficient identification of novel gene–disease associations. The interpretation of WGS/WES data linked to individuals is increasingly being used to identify causal variants that may lead to an abnormal phenotype or a disease Krier *et al*. (2016). Despite its success, these approaches leave many patients undiagnosed, with estimated diagnostic yields of 25%-50% Clark *et al*. (2018).

While there have been several efforts to predict and prioritize pathogenic genomic variants, in particular, Single Nucleotide Polymorphisms (SNPs) and small Insertion or Deletion (InDels) Eilbeck *et al*. (2017), predicting the functional impact of variants discovered through genome sequencing studies remains challenging. This is due to the limited gene–phenotype information available; also, variants may cover multiple coding, non-coding, or intergenic regions and overlap several genes Shameer *et al*. (2016). Existing methods for predicting the pathogenicity of genomic variants may be based on the impact of variants on protein structure, measures of sequence conservation, or function by relying only on the genomic sequence information Eilbeck *et al*. (2017). While several methods exist to identify disease-associated variants in patient cohorts, it is more challenging to discover disease-associated variants that exist in a single sample or pedigree, in particular in rare Mendelian disorders Sanchis-Juan *et al*. (2018).

Another group of methods for finding variants causing abnormal phenotypes predicts variant pathogenicity and prioritize damaging variants using the relation between the phenotypes of a patient and the phenotypes in a database of genotype–phenotype associations Köhler *et al*. (2014). Phenotype-driven variant prioritization methods aim to link variants to the phenotypes observed in individuals using prior knowledge Eilbeck *et al*. (2017). Commonly, the link is established using a similarity measure between phenotypes associated with a variant or gene and the phenotypes observed in a patient Smedley *et al*. (2015). Phenotype-based methods are successful in finding disease-associated variants Shefchek *et al*. (2020a) but suffer from the limited information about variant– or gene–phenotype associations. One way to overcome this limitation is to utilize and link the phenotypes observed in model organisms to human phenotypes Shefchek *et al*. (2020a). However, even when including phenotypes from model organisms, a large number of human protein-coding genes remain without associations, thereby limiting the success of phenotype-based methods to variants or genes that have previously been studied either in human or animal models or relying on guilt-by-association approaches in which information about phenotypes is propagated through associations such as interaction networks Smedley *et al*. (2014).

Several deep learning and machine learning methods are now available that can predict phenotypes from genotype Zhou *et al*. (2019); Kulmanov and Hoehndorf (2020) or associate phenotypes with different types of information available for genes, including the functions of gene products and anatomical sites of expression Chen *et al*. (2020); Smaili *et al*. (2019). These methods use machine learning to relate information through background knowledge contained in formalized knowledge bases, or ontologies, and can accurately identify phenotype-associated genes without prior knowledge about phenotypes, often significantly improving over the use of semantic similarity measures Kulmanov *et al*. (2020). A limitation of these methods is that they are usually transductive instead of inductive Kulmanov *et al*. (2020), i.e., the diseases or disorders for which associated genes are predicted should already be available at the time of training the model. As these methods require information about disease-associated phenotypes during training, they cannot generalize to entirely new cases, thereby limiting their application in identifying phenotype-associated genomic variants. Another limitation can be biases introduced by the neural network and the phenotypes annotations Alghamdi *et al*. (2022) or similarity measure Kulmanov and Hoehndorf (2017).

We developed Embedding Pathogenicity Variant Predictor (EmbedPVP), a computational method to prioritize variants that are pathogenic and involved in the development of specific phenotypes or genetic diseases. EmbedPVP prioritizes single nucleotide variants or small insertions or deletions involved in genetic diseases. Our method combines genomic information and clinical phenotypes and leverages a large knowledge base derived from human and model organisms for knowledge-enhanced learning. We use different neuro-symbolic embedding-based methods to learn from the background knowledge and combine the information from embedding and pathogenicity prediction to predict the variant that most likely causes the phenotypes observed in the patients. We demonstrate that our method improves over the state-of-the-art in detecting disease-associated variants in multiple benchmark datasets. We have made EmbedPVP freely available as a Python package at https://github.com/bio-ontology-research-group/EmbedPVP.

## 2 Materials and Methods

### 2.1 Genotype and clinical phenotype datasets

We performed all of our experiments on a set of pathogenic and disease-causing variants for diseases collected from different databases. We inserted the variants we obtained into synthetic genomes with a set of benign, pathogenic, and unknown variants from the 1,000 Genomes Project Consortium *et al*. (2012). We use three different datasets of variants to generate synthetic patients and evaluate the performance of EmbedPVP. The first dataset, the PAVS-synthetic dataset, covers clinically validated Saudi variants from an in-house database, the Phenotype-Associated Variants in Saudi Arabia (PAVS) database Ali Raza Syed and Althagafi (2022) representing 1528 individuals. PAVS is a database that combines a set of clinically validated pathogenic variants with a set of manually curated pathogenic variants observed in the genomes of the Saudi population and their associated phenotypes. All phenotypes are mapped to their HPO identifiers. The second dataset, Phenopackets Jacobsen *et al*. (2022), represents 384 individuals described in published case reports with HPO terms and their causal genetic variants. As the final dataset, we selected 1082 newly inserted pathogenic variants (between 2022-01-04 and 2022-10-31) from the ClinVar database Landrum *et al*. (2020). We further subsetted these datasets to cover other evaluations, such as exonic vs. non-exonic variants (Supplementary Materials Section 2.2), variants in overlapping and intergenic regions (Supplementary Section 2.3), variants in genes with no phenotype annotations (Supplementary Section 2.4), newly discovered genes, and diseases observed or not observed during the training.

### 2.2 Resources for ontologies and annotation phenotypes

We use four primary ontologies: Human Phenotype Ontology (HPO), Mammalian Phenotype Ontology (MP), Gene Ontology (GO), and the Uberon cross-species integrated anatomy ontology (UBERON).

First, the phenotypes associated with human genes were downloaded from the HPO database on May 30, 2022. We obtained the phenotype annotations for 4318 human genes, including 205, 429 associations between genes and HPO. Second, the phenotypes associated with mouse genes and the orthologous gene mappings from mouse genes to human genes were obtained from the Mouse Genome Informatics (MGI) database Smith and Eppig (2009), downloaded on June 7, 2022. We obtained phenotype annotations for 13, 529 mouse genes, including 228, 214 associations between genes and MP classes. We mapped each mouse gene to its human ortholog using the file `HMD_HumanPhenotype.rpt` available at the MGI database, resulting in 9, 879 human genes for which the mouse ortholog has phenotype associations. Third, we used biological function (GO) annotations from the GO website Ashburner *et al*. (2000) downloaded on March 14, 2022. We collected 18, 495 human gene products (495,719 annotations in total). We mapped the UniProt accessions to Entrez gene identifiers using the mappings provided by the Entrez database Maglott *et al*. (2010), and we obtained 17,786 Entrez genes for which the gene product has GO annotations. Fourth, for the anatomical location of gene expression, we downloaded the Tissue Expression Profiles (GTEx) dataset GTEx Consortium (2015) from the Gene Expression Atlas Papatheodorou *et al*. (2020), which characterizes gene expression across 53 tissues. We mapped the Ensembl protein identifiers to Entrez gene identifiers using the mapping provided by the Entrez database Maglott *et al*. (2010). We obtained 20, 538 Entrez genes, which have expression levels above the 4.0 threshold in one or more tissue. We mapped each tissue to the UBERON ontology, excluding the expression in *EBV-transformed lymphocyte* and *transformed skin fibroblast* since these two tissues are not available in the UBERON ontology.

Finally, because these annotations are available for different numbers of genes, we also used the phenotypes based on the union of all genes and their annotations (i.e., for genes that have annotations from one, two or all four datasets, HPO, MP, GO, and Uberon). We used the integrated phenotype ontology uPheno Shefchek *et al*. (2020b) as our phenotype ontology to add background knowledge from biomedical ontologies, as it integrates human and model organism phenotypes and allows them to be compared.

To evaluate gene–disease associations, we used the phenotypes available in the HPO database Köhler *et al*. (2019) to associate diseases from the Online Mendelian Inheritance in Men (OMIM) database Amberger *et al*. (2011) to their phenotypes. In total, we have 4431 OMIM diseases and 3418 genes in our knowledge base, representing 7405 associations; we used 80% of these associations during supervised training to generate the representations, 15% for the validation, and 5% for testing.

### 2.3 Generation of synthetic patients and synthetic phenotypes

We created synthetic genotypes in Variant Call Format (VCF) format using the reference genome from the 1000 Genomes Project. We simulated a more realistic genome by randomly selecting 100,000 variants such that 90% are in intronic regions and 10% as exonic within regions. We then filtered the variants to select variants with MAF *<*1% in 1000 Genomes Project, ExAC Karczewski *et al*. (2017), and gnomAD Karczewski and Francioli (2017) databases for all the population, and as a result, we obtained 98, 194 variants to represent our synthetic genome. We then inserted the causative pathogenic variants from our evaluation cohorts (PAVS, Phenopackets, ClinVar Time-based split) into the synthetic genome, which, together with the associated phenotypes, represents the synthetic patients.

For the phenotypes linked to each patient, we evaluated the phenotypes reported for each patient (using PAVS and Phenopackets cohorts). In addition, since we have the OMIM diseases reported for each causative variant, we performed the same experiments using the phenotypes linked with the disease in HPO, which represents more phenotypic variability compared to the reported phenotypes. We used the VCF files together with the HPO phenotypes, either clinical or from OMIM, to run the different models. We then ranked the inserted variants using EmbedPVP models and other prioritization methods.

### 2.4 Generation of ontology annotation-based embeddings

Formally, we define an ontology using a signature Σ = (**C, R, I**), where **C, R, I** are sets corresponding to ontology concepts, roles, and individuals, respectively. An embedding is a structure-preserving mapping between two mathematical structures. To generate embeddings from ontology entities into a vector representation, we followed different approaches identified and categorized in Kulmanov *et al*. (2020) such as (1) graph-embeddings with random walks, (2) graph embeddings with knowledge graph embeddings methods, and (3) model-theoretic embeddings. To predict gene–disease associations, we used a scoring function *s* given by the embedding method. For a gene *g* and a disease *d, s*(*g, d*) will output a value in the range [0, 1] indicating the plausibility of the association to hold true. The first and second approaches require the generation of a graph out of the ontology axioms. This process is called *graph projection* Zhapa-Camacho and Hoehndorf (2023). The graph projection methods we chose are the one found in DL2Vec Chen *et al*. (2020) and OWL2Vec* Chen *et al*. (2021).

A relational graph is a tuple *G* = (*V, E, L*) with sets *V* of vertices, *L* of edge labels and *E* ⊆ *V* × *L* × *V* of edges represented as triples (*h, r, t*) where *h, t* are nodes and *r* is an edge label. Given a graph, a random walk *w* = {*v*_0_, *v*_1_, *v*_2_, …, *v*_*n*_} of length *n* is constructed iteratively by choosing an initial node *v*_0_ ∈ *V* and obtaining nodes *v*_*i*+1_ = *next*(*v*_*i*_) given by the function *next*. After generating a graph using the projection function in DL2Vec, we used DeepWalk Perozzi *et al*. (2014) to create *k* walks of size *n* for each node in the graph. Traditionally, in DeepWalk, the *next*(*v*_*i*_) function generates the element *v*_*i*+1_ by choosing randomly from the neighbors of *v*_*i*_. However, to include edge label information, we used a variation from DeepWalk that takes not only neighboring vertices but also the edge label between them. Thus, a walk with size *n* will contain 2*n* − 1 elements.

To capture the co-occurrence of ontology entities, we trained a Word2Vec model, where the input is the collection of *k* · |*V* | random walks. Word2Vec, under the Skip-gram architecture, aims to find word representations that are useful to predict surrounding words Mikolov *et al*. (2013). Thus, given a sequence *v*_0_, *v*_1_, …, *v*_*n*_, the training objective is 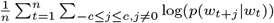 where *p* is the softmax function. Given that Word2Vec can capture the co-occurrence of entities, we chose a similarity-based scoring function defined as 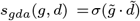 where 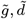 are the vector representations obtained by training the Word2Vec model, (·) correspond to the dot product and *σ* is the sigmoid function.

Graph embeddings using random walks generate embeddings that are useful for computing similarity between nodes, but they do not consider the relation during the similarity computation. To incorporate relations, we generated embeddings by using Knowledge Graph Embedding methods (KGE) Wang *et al*. (2017), such as TransE Bordes *et al*. (2013), TransR Lin *et al*. (2015), TransD Ji *et al*. (2015), DistMult Yang *et al*. (2014), and ConvE Dettmers *et al*. (2018). All KGE methods implement a scoring function *s*(*h, r, t*) indicating the plausibility of the triple (*h, r, t*) to exist in the graph. The training objective is 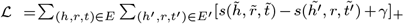 where the set *E*^’^ is the set of negative triples (i.e., triples not existing in the graph) generated by either corrupting the head or tail of a positive triple in *E*. The training objective aims to minimize the score of a positive triple and maximize the scores of negative triples.

To predict gene–disease associations for a gene *g* and a disease *d*, we compute *s*_*gda*_(*g, d*) = *s*(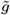, is_associated_with, 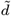).

Graph-based methods ignore semantic information of ontology axioms. Concept descriptions **C** in the Description Logic εℒ can be constructed as any of the normal forms *C* ⊑ *D,C* ⊓ *D* ⊑ *E, C* ⊑ ∃*R*.*C* and ∃*R*.*C* ⊑ *D*. An interpretation ℐ = (Δ^ℐ^, ·^ℐ^) is given by an non-empty domain Δ^ℐ^ and an interpretation function mapping every concept *C* ∈ **C** to a set *C*^ℐ^ ⊆ Δ^ℐ^ and every role *R* ∈ **R** to a set *R*^ℐ^ ⊆ Δ^ℐ^ × Δ^ℐ^. Moreover, the interpretation function maps complex concept descriptions as follows: ⊥^ℐ^ = Ø, T^ℐ^= Δ^ℐ^, (*C* ⊓ *D*) ^ℐ^ = *C*^ℐ^ ∩ *D*^ℐ^, (∃*R*.*C*) ^ℐ^ = {*a* ∈ Δ^ℐ^ | ∃*b* ∈ Δ^ℐ^ : (*a, b*) ∈ *R*^ℐ^ ∧ *b* ∈ *C*^ℐ^} An interpretation ℐ is a *model* if for every axiom *C* ⊑ *D* the inclusion *C*^ℐ^ ⊆ *D*^ℐ^ holds.

In order to incorporate semantic information, we used two geometric-based embedding methods: ELEmbeddings Kulmanov *et al*. (2019) and ELBoxEmbeddings Peng *et al*. (2022). These methods represent ontology concepts as geometric bodies such as *n*-dimensional balls and *n*-dimensional boxes, respectively. For every axiom *C* ⊑ *D*, the training objective minimizes the inclusion loss of the geometric representation of *C* within the geometric representation of *D*. Therefore, the scoring method for every axiom is *s*(*C* ⊑ *D*) = *inclusion*(*C, D*).

The *inclussion* function is defined for each normal form in ELEmbeddings and ELBoxEmbeddings. The training objective follows a similar approach as in Eq 3, where the positive samples are the axioms in the ontology, and the negative samples are generated by corrupting the concept names on the right-hand side of the axiom. To predict gene–disease associations, we compute the score of the axiom: 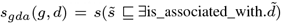.

### 2.5 Updating embedding models to handle new phenotypes

In order to assess the generalizability of the models, we further evaluate the patients using an inductive approach. To achieve this, we first trained each model with an initial set of diseases from OMIM until a convergence criterion is reached. The resulting trained models, along with their corresponding embeddings, were then saved for subsequent inductive utilization. Following this, we added information of the phenotypes of each new disease *d*^’^ phenotypes separately into the model. Subsequently, the model was trained for a small number of iterations to update the embedding representations with the new set of phenotypes. Supplementary Algorithm 1, showed the details of updating the trained model.

Using an inductive setting enables the insertion of new patient information with low computational cost because the previously learned knowledge can be reused to constrain the new knowledge. Compared to a transductive approach, where adding new information requires training a model from scratch, inductive learning allows reusability of the trained models, therefore enabling better scalability properties than a transductive approach.

We generated the embedding representation with transductive and inductive settings using different embedding-based approaches implemented by the mOWL library Zhapa-Camacho *et al*. (2023). We used different parameter sets according to each method. In the Supplementary Section 1, we report the different tuned parameters used for each embedding method.

### 2.6 Functional variant feature

We annotate variants with a set of genomic features using public databases. We use Annovar Wang *et al*. (2010), which uses data from multiple external databases. From the annotations provided by Annovar, we use the type of variants and the gene information. While not used as a feature of our prediction model, we also use Annnovar to identify the allele frequency of variants using the 1000 Genomes allele frequency Sudmant *et al*. (2015) and allele frequency from gnomAD Collins *et al*. (2020). We use this information to filter out common variants before applying our predictions. For the pathogenicity prediction, we rely on the Combined Annotation Dependent Depletion (CADD) Rentzsch *et al*. (2019) score. CADD is a tool for scoring the deleteriousness of single nucleotide variants and insertion/deletion variants in the human genome.

### 2.7 Estimating variant pathogenicity

We hypothesize that the phenotypes we observe in patients that result from a variant correlates with the phenotypes observed when altering one or more of the genes affected by the variant and, therefore, that phenotypes associated with genes can provide additional information to predict potential causative variants Doelken *et al*. (2012). We combine the score from the embedding method we employ with the CADD pathogenicity score to provide the final prediction.

### 2.8 Performance evaluation and comparison

We compare EmbedPVP with variant prioritization tools based on the genotype information, specifically CADD Kleinert and Kircher (2021), SIFT Ng and Henikoff (2003), PolyPhen2 Adzhubei *et al*. (2013), MetaSVM Sun and Yu (2019), and DANN Quang *et al*. (2015). We also evaluated different phenotype-based methods, PhenIX Zemojtel *et al*. (2014), Exomiser-hiPHIVE Robinson *et al*. (2014), PHIVE Robinson *et al*. (2014), and DeepPVP Boudellioua *et al*. (2019). We assessed their effectiveness in the different benchmark datasets. We evaluated the performance of our models and baseline methods by calculating the recall at different ranks, i.e., finding the rank of the inserted variants and then reporting the top hits, top 10, top 30, and top 50 hits.

## 3 Results

### 3.1 Overview of the EmbedPVP model

The EmbedPVP workflow consists of a hierarchical architecture that utilizes a set of SNPs or InDels in VCF format as input, along with a set of phenotypes encoded using HPO. EmbedPVP generates a ranked list of variants from the input VCF file based on their probability of being associated with the input set of phenotypes. EmbedPVP utilizes a knowledge base with different ontologies and their annotations (Supplementary Figure 1.A) to relate genotypes to phenotype, EmbedPVP generates embedding representations for the input phenotypes and genes using embedding methods (Figure 1.B). Next, for the input set of variants, EmbedPVP collects genomic features and retrieves for each variant the gene or genes to which it is related; coding variants are related to the gene in whose coding region they lie, and intergenic variants are related to their closest genes. If a variant lies in the coding region of multiple genes, it relates to each of them (Figure 1.C). EmbedPVP then calculates the similarity between the input set of phenotypes and the ontology-based embedding for the genes using the scoring function associated with the selected embedding method. Finally, EmbedPVP uses the weighted averages of the similarity score with the pathogenicity prediction as the final prediction score for the variant (Figure 1.C).

### 3.2 EmbedPVP evaluation

We conducted evaluations on various benchmark datasets, including synthetic datasets using clinical phenotypes and OMIM phenotypes. For this purpose, we trained EmbedPVP with a unique representation for each sample based on its phenotypes, i.e., all samples were already known during embedding generation. In other words, we perform transductive inference. The evaluations aimed to assess the performance of different embedding methods on these datasets. Table 1 provides a comparison of the performance of EmbedPVP against other state-of-the-art methods using the PAVS dataset (refer to Supplementary Table 1 for the results of all other methods). Additionally, Supplementary Figure 2 shows the average ranks for hits at 1, 10, 30, and 50.

**Table 1.**
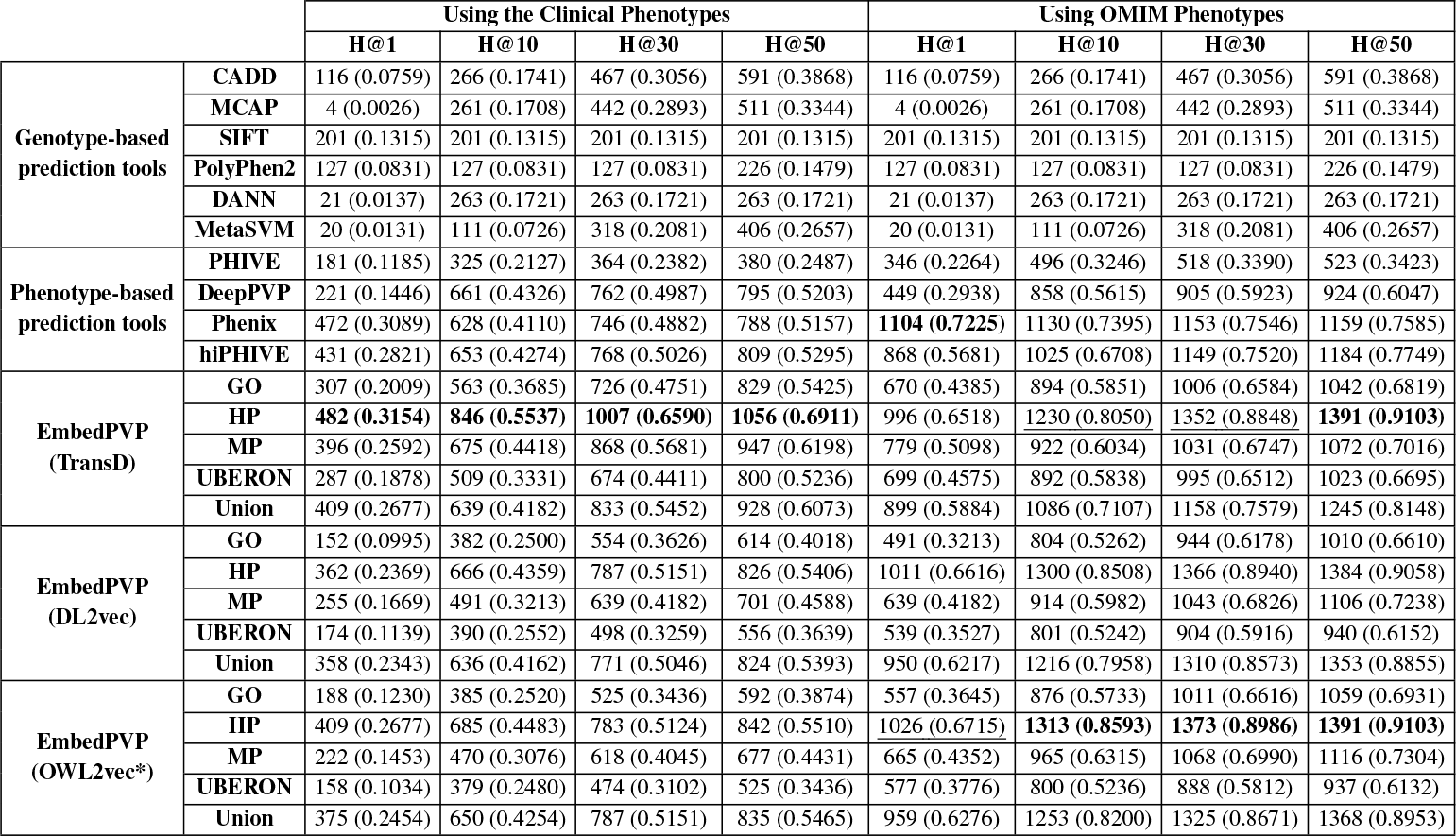
EmbedPVP variant prediction results across several ontologies with different neuro-symbolic knowledge embedding methods.

Based on the results, we observed that EmbedPVP using the TransD model with HP ontology achieved the highest performance among the phenotype-based prediction tools using clinical phenotypes. However, when using OMIM phenotypes, OWL2vec* demonstrated slightly better performance. DL2vec and OWL2vec* performed similarly in both clinical and OMIM phenotypes compared to other phenotype-based models. These findings suggest that TransD captures the complex representations of relationships more effectively. TransD utilizes a translation-based approach to model relationships, which enables it to capture multi-relational relationships between entities. On the other hand, OWL2vec* and DL2vec primarily focus on representing hierarchical relationships using the ontology’s structure. Although they excel at capturing hierarchical relationships, they may struggle to represent more intricate relationships involving multiple entities or more complex representations, in contrast to the TransD model.

When evaluating the Phenopackets dataset (Supplementary Table 3), we observed that the EmbedPVP (DL2vec) method performed better in terms of top hits. However, among the phenotype-based methods, Phenix demonstrated better performance for the remaining metrics.

Furthermore, we evaluated our method using ClinVar time-split variants, and the results of the different methods are presented in Supplementary Table 4. In this dataset, EmbedPVP using the TransD method outperforms other methods using the HP model for the top hits, the Union model for the H@10, and the GO model for H@30 and H@50. To further assess the model’s performance and remove potential biases due to partial information about gene–disease tuples being present during training, we conducted additional evaluations by splitting the dataset based on novel genes and diseases that were not present during training. We created different subsets, as follows: A. novel genes and diseases (454 variants), B. novel genes and known diseases (31 variants), C. novel diseases and known genes (111 variants), and D. known genes and diseases (484 variants). The results for these different subsets are shown in Supplementary Table 5 for A and B, and Table 6 for C and D. We also noticed the EmbedPVP models performed better compared to the other phenotype-based and genotype-based methods.

### 3.3 Improved generalization to new phenotypes with inductive inferences

We evaluate the performance of inductive inference using PAVS with clinical phenotypes, with selected models based on the best-performing transductive approach, including the OWL2Vec*, DL2vec, TransE, and TransD embedding models. Supplementary Table 10 presents the results comparing the inductive and transductive approaches. The results show a drop in performance, and as a consequence, the inductive model does not perform better than other methods. This result demonstrates that the additional time required for retraining EmbedPVP in the presence of new individuals to analyze is necessary for its performance.

## 4 Discussion

We developed a method for prioritizing candidate causative variants when given a set of disease-associated phenotypes and genotypes. Our approach utilizes various features characterized through ontologies and employs neuro-symbolic embedding methods to exploit the information in ontologies and their annotations. As a result, EmbedPVP can improve phenotype-based prediction of disease-causing variants. Moreover, we also explored the impact of clinical phenotype descriptions and could demonstrate that the embeddings we utilize are robust to noisy phenotype descriptions.

Knowledge-enhanced learning involves the utilization of background knowledge to enhance predictive models. Knowledge-enhanced learning is especially useful when too little training data is available to apply supervised learning directly, and where structured knowledge is available that can constrain search Feigenbaum *et al*. (1977). The large number of biomedical ontologies and the knowledge they contain has been used deductively to generate additional knowledge that could then be used to improve machine learning tasks Köhler *et al*. (2013); Matentzoglu *et al*. (2019); Hoehndorf *et al*. (2011); in our work, we use the background knowledge in ontologies not deductively but rather as part of a neuro-symbolic method Hitzler and Sarker (2022) where a form of inference happens in a latent space Shakarian *et al*. (2023); Hitzler *et al*. (2023).

In our application, we rely on axioms from the Gene Ontology Ashburner *et al*. (2000), phenotype ontologies, and anatomy ontologiesSmith *et al*. (2007). We use these ontologies to integrate information about pathways, interactions between genes, anatomical site of gene expression, and protein functions, and ontologies already link all this information to phenotypes using formal axioms. In particular, phenotype ontologies have long been constructed using the entity-quality (EQ) method where phenotypes are decomposed into an affected entity (an anatomical site, or a biological function) and a quality (using the PATO ontology of qualities) Gkoutos *et al*. (2018); Mungall *et al*. (2010). Using these axioms now proves useful not only for data integration (which was one of the original intentions in developing these axioms) but also enables knowledge-enhanced learning in these domains.

EmbedPVP is not the first approach that uses ontology semantics in detecting genotype–phenotype relations; in particular semantic similarity measures have been used for a long time to predict gene–disease associations Köhler *et al*. (2009), and semantic similarity measures have also been incorporated in variant prioritization tools such as Exomiser Robinson *et al*. (2014). While semantic similarity measures are able to compare sets of classes from a single ontology, our neuro-symbolic approach is able to “learn” a similarity measure within a latent space, and determine the similarity between classes that are related through complex and heterogeneous axioms. This property allows us not only to improve predictive performance over approaches that rely on semantic similarity (such as the Exomiser tool), but, maybe more importantly, extends the scope of phenotype-similarity methods for finding candidate disease genes to genes for which no phenotypes are known. Previously, a major advance has been the use of model organism phenotypes to expand the scope of methods that find disease-associated genes or variants through comparison to patient phenotypes Chen *et al*. (2012); Hoehndorf *et al*. (2011); the combination of mouse and zebrafish phenotypes spans a large part of the human genome, but still there are gaps where no phenotypes are associated with a gene. EmbedPVP can apply phenotype similarity for any gene for which a site of expression or gene function is known.

One main limitation of EmbedPVP is that it uses a transductive method which requires retraining parts of the model when a new case or set of cases is analyzed. This is mainly a limitation of time as re-training is part of applying EmbedPVP to a new case; however, in particular, when analyzing larger number of cases, it may still be reasonable to retrain and then predict. In the future, however, we intend to focus our efforts on designing novel strategies for inductive inference.

## 5 Conclusion

We developed EmbedPVP, a method for prioritizing candidate causative variants given a set of abnormal phenotypes. Our method applies machine learning to background knowledge integrated through through ontologies and not only improves the phenotype-based prediction of disease-associated variants, but also extends phenotype-based variant prioritization to variants in genes for which no phenotypes are available; instead, EmbedPVP can use knowledge about gene functions, sites of expression, interactions, and also phenotypes in humans or model organisms to prioritize variants. We implemented and evaluated different embedding-based methods for learning from biomedical knowledge bases, applying graph-based as well as model-based methods. EmbedPVP is an end-to-end model and is applicable not only to single nucleotide variants in coding regions, but also to non-coding variants and small insertions and deletions. EmbedPVP has been designed to prioritize variants even when phenotype information is missing or noisy, and EmbedPVP could improve the prediction of causative variants even in the presence of noise. EmbedPVP improves over state-of-the-art methods for phenotype-based variant prioritization, particularly in improving the recall in finding phenotype-associated variants across various benchmark datasets. EmbedPVP is freely available at https://github.com/bio-ontology-research-group/EmbedPVP.

## Acknowledgements

We acknowledge the use of computational resources from the KAUST Supercomputing Core Laboratory.

