## Supplementary materials for "Prioritizing genomic variants through neuro-symbolic, knowledge-enhanced learning"

#### 1 Hyperparameter

This supplementary material presents the results of different embedding methods across various categories. We have classified them into three groups: graph–based methods, translation embedding methods, and semantic methods. For each category, we have evaluated different sets of parameters as follows:

- **Graph–based methods:**
  - vector_size: [50, 100, 200]
  - window: [5, 10, 20]
  - epochs: [10,20, 30, 50, 100]
  - num_walks: [10, 30, 50, 80]
  - walk_length: [5, 10, 20, 30]
- **Translation-based methods:**
  - vector_size: [50, 100, 150, 200]
  - epochs: [20, 50, 100, 150, 200]
  - learning_rate: [0.0001, 0.001, 0.01, 0.1]
  - batch_size: [2048, 4096, 8192]
- **Semantic-based methods:**
  - vector_size: [50, 100, 150]
  - margin: [0.1, 0.2,0.01]
  - neg_norm: [1,0.5]
  - epochs: [20, 100, 1000]
  - learning_rate: [0.0001, 0.001, 0.01]
  - batch_size: [2048, 4096, 8192]

**Algorithm 1** Incorporating New Patient Information and Updating Embeddings

**Input:** Description Logic theory $D$ with signature $\Sigma = (\mathbf{C}, \mathbf{R}, \mathbf{I})$ in ontology OWL format, Trained Embeddings $E_{trained}$ for $\mathbf{C}$ and $\mathbf{R}$, New Patient Information $P = \{patient_i \sqsubseteq \exists has\_phenotype.phenotype_j | \text{ for } i \in [1, n] \text{ for } j \in [1, m]\}$ as DL axioms
**Output:** Updated Knowledge Graph $G' = (V', E')$, Updated Embeddings $E_{\text{updated}}$
**procedure** IncorporateAndUpdate($D, E_{\text{trained}}, P$)
    Generate $D'$ with signature $\Sigma' = (\mathbf{C}', \mathbf{R}', \mathbf{I})$ by adding axioms $P$ to $D$
    Generate new graph $G$ from $D'$
    $new\_nodes \leftarrow \mathbf{C}'/\mathbf{C}$
    $corpus \leftarrow$ Random walks on the updated knowledge graph $G' = (V', E')$ start from nodes in $new\_nodes$
    Initialize Word2Vec model with $E_{trained}$ embeddings and add random embeddings for entities in $new\_nodes$
    Train Word2Vec embeddings on $corpus$
    $E_{\text{updated}} \leftarrow$ Updated embeddings from training on $G'$
    **return** $G' = (V', E')$, $E_{\text{updated}}$
**end procedure**

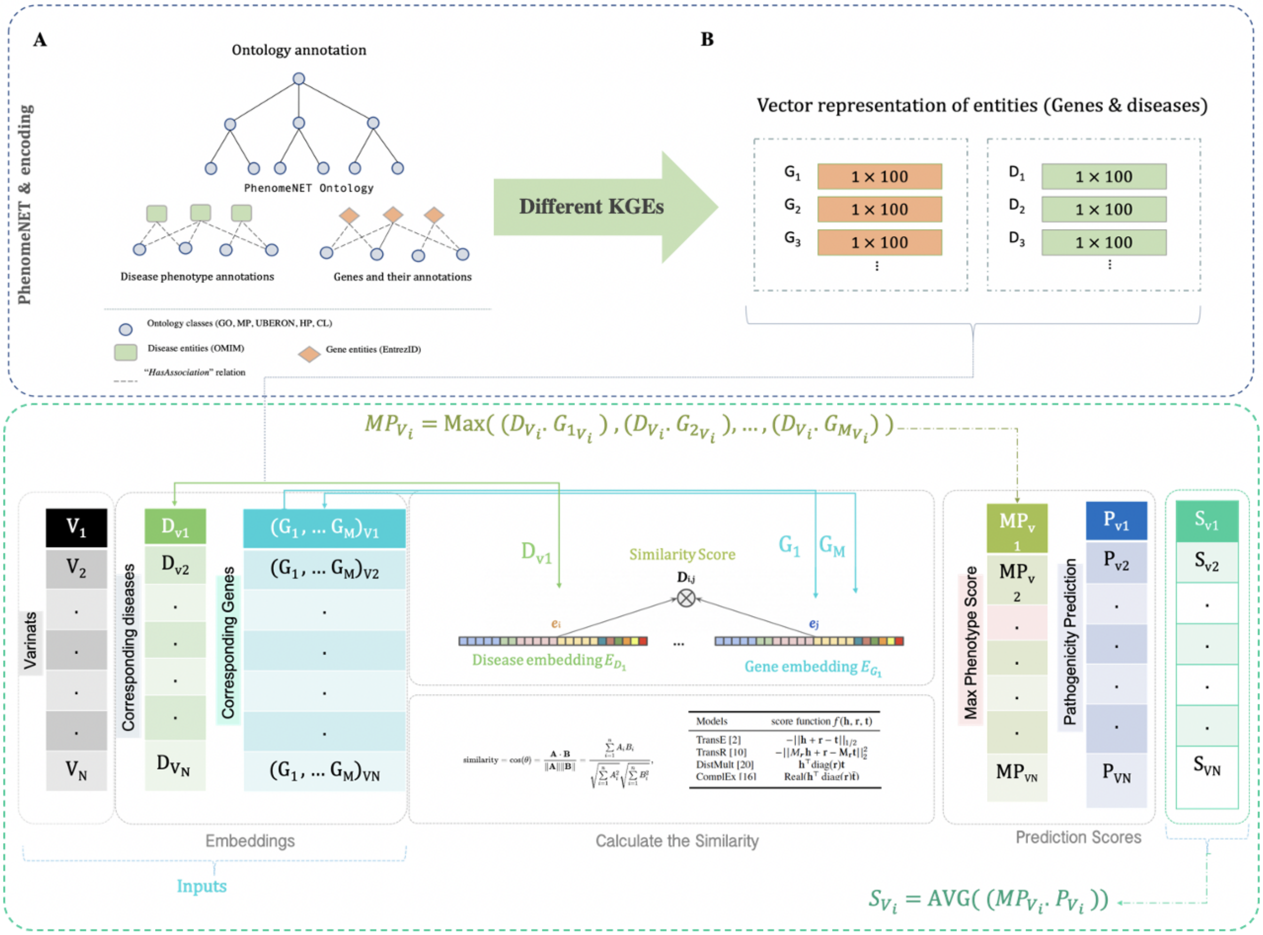

Figure 1: Model workflow for EmbedPVP. **A)** EmbedPVP first generates the background knowledge with different ontologies annotations. **B)** Generates embedding representation for the disease and genes using different embedding methods. **C)** Calculates the similarity for the gene-disease pairs for the embedding representation and averages the similarity score with pathogenicity prediction.

### 2 Variant predictions results across several models

We have conducted evaluations on various benchmark datasets, including synthetic datasets using clinical phenotypes and OMIM phenotypes, using the `transductive approach`. The evaluation assesses the performance of different embedding methods on these different datasets. The following tables and figures present a comparison of their performance with other state-of-the-art methods.

#### 2.1 Transductive approach

##### 2.1.1 PAVS dataset

Table 1, Figure 2, or Figure 3, for the exonic variants and other types in Table 2

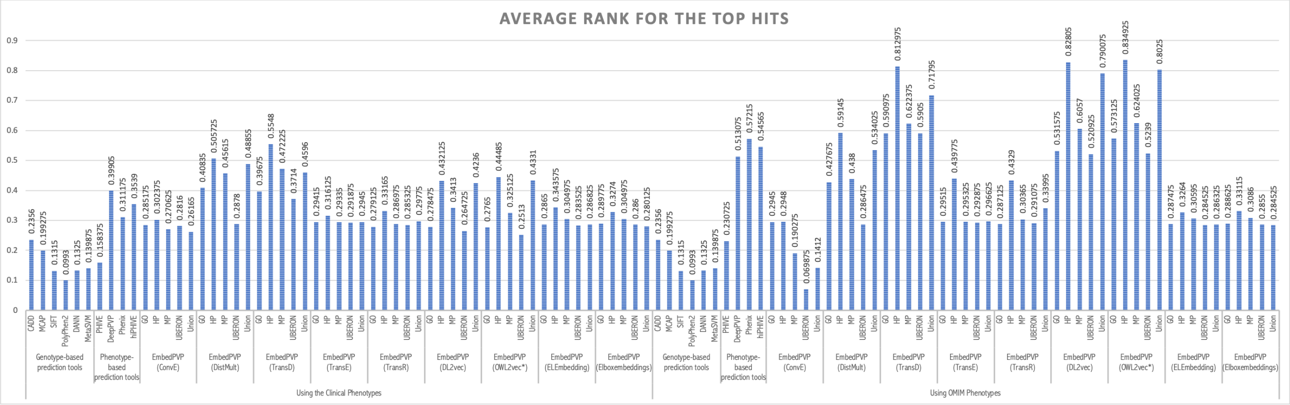


Figure 2: Average ranks for recall at 1, 10, 30, and 50 using synthetic datasets with clinical phenotypes and OMIM phenotypes (PAVS datasets)

| | | Using the Clinical Phenotypes | | | | Using OMIM Phenotypes | | | |
|---|---|---|---|---|---|---|---|---|---|
| | | **H@1** | **H@10** | **H@30** | **H@50** | **H@1** | **H@10** | **H@30** | **H@50** |
| **Genotype-based prediction tools** | **CADD** | 116 (0.0759) | 266 (0.1741) | 467 (0.3056) | 591 (0.3868) | 116 (0.0759) | 266 (0.1741) | 467 (0.3056) | 591 (0.3868) |
| | **MCAP** | 4 (0.0026) | 261 (0.1708) | 442 (0.2893) | 511 (0.3344) | 4 (0.0026) | 261 (0.1708) | 442 (0.2893) | 511 (0.3344) |
| | **SIFT** | 201 (0.1315) | 201 (0.1315) | 201 (0.1315) | 201 (0.1315) | 201 (0.1315) | 201 (0.1315) | 201 (0.1315) | 201 (0.1315) |
| | **PolyPhen2** | 127 (0.0831) | 127 (0.0831) | 127 (0.0831) | 226 (0.1479) | 127 (0.0831) | 127 (0.0831) | 127 (0.0831) | 226 (0.1479) |
| | **DANN** | 21 (0.0137) | 263 (0.1721) | 263 (0.1721) | 263 (0.1721) | 21 (0.0137) | 263 (0.1721) | 263 (0.1721) | 263 (0.1721) |
| | **MetaSVM** | 20 (0.0131) | 111 (0.0726) | 318 (0.2081) | 406 (0.2657) | 20 (0.0131) | 111 (0.0726) | 318 (0.2081) | 406 (0.2657) |
| **Phenotype-based prediction tools** | **PHIVE** | 181 (0.1185) | 325 (0.2127) | 364 (0.2382) | 380 (0.2487) | 346 (0.2264) | 496 (0.3246) | 518 (0.3390) | 523 (0.3423) |
| | **DeepPVP** | 221 (0.1446) | 661 (0.4326) | 762 (0.4987) | 795 (0.5203) | 449 (0.2938) | 858 (0.5615) | 905 (0.5923) | 924 (0.6047) |
| | **Phenix** | 472 (0.3089) | 628 (0.4110) | 746 (0.4882) | 788 (0.5157) | **1104 (0.7225)** | 1130 (0.7395) | 1153 (0.7546) | 1159 (0.7585) |
| | **hiPHIVE** | 431 (0.2821) | 653 (0.4274) | 768 (0.5026) | 809 (0.5295) | 868 (0.5681) | 1025 (0.6708) | 1149 (0.7520) | 1184 (0.7749) |
| **EmbedPVP (ConvE)** | **GO** | 195 (0.1276) | 366 (0.2395) | 523 (0.3423) | 659 (0.4313) | 162 (0.1060) | 389 (0.2546) | 567 (0.3711) | 682 (0.4463) |
| | **HP** | 192 (0.1257) | 389 (0.2546) | 558 (0.3652) | 709 (0.4640) | 201 (0.1315) | 388 (0.2539) | 535 (0.3501) | 678 (0.4437) |
| | **MP** | 154 (0.1008) | 385 (0.2520) | 514 (0.3364) | 601 (0.3933) | 97 (0.0635) | 232 (0.1518) | 381 (0.2493) | 453 (0.2965) |
| | **UBERON** | 183 (0.1198) | 351 (0.2297) | 521 (0.3410) | 666 (0.4359) | 35 (0.0229) | 80 (0.0524) | 139 (0.0910) | 173 (0.1132) |
| | **Union** | 159 (0.1041) | 315 (0.2062) | 494 (0.3233) | 631 (0.4130) | 71 (0.0465) | 158 (0.1034) | 270 (0.1767) | 364 (0.2382) |
| **EmbedPVP (DistMult)** | **GO** | 322 (0.2107) | 564 (0.3691) | 761 (0.4980) | 849 (0.5556) | 305 (0.1996) | 608 (0.3979) | 817 (0.5347) | 884 (0.5785) |
| | **HP** | 397 (0.2598) | 808 (0.5288) | 918 (0.6008) | 968 (0.6335) | 552 (0.3613) | 976 (0.6387) | 1029 (0.6734) | 1058 (0.6924) |
| | **MP** | 364 (0.2382) | 645 (0.4221) | 841 (0.5504) | 938 (0.6139) | 347 (0.2271) | 595 (0.3894) | 821 (0.5373) | 914 (0.5982) |
| | **UBERON** | 148 (0.0969) | 361 (0.2363) | 570 (0.3730) | 680 (0.4450) | 157 (0.1027) | 373 (0.2441) | 551 (0.3606) | 670 (0.4385) |
| | **Union** | 352 (0.2304) | 729 (0.4771) | 915 (0.5988) | 990 (0.6479) | 418 (0.2736) | 840 (0.5497) | 985 (0.6446) | 1021 (0.6682) |
| **EmbedPVP (ELEmbedding)** | **GO** | 183 (0.1198) | 340 (0.2225) | 540 (0.3534) | 688 (0.4503) | 196 (0.1283) | 341 (0.2232) | 540 (0.3534) | 680 (0.4450) |
| | **HP** | 203 (0.1329) | 458 (0.2997) | 649 (0.4247) | 790 (0.5170) | 179 (0.1171) | 432 (0.2827) | 639 (0.4182) | 745 (0.4876) |
| | **MP** | 184 (0.1204) | 404 (0.2644) | 549 (0.3593) | 727 (0.4758) | 180 (0.1178) | 426 (0.2788) | 536 (0.3508) | 728 (0.4764) |
| | **UBERON** | 179 (0.1171) | 342 (0.2238) | 534 (0.3495) | 678 (0.4437) | 182 (0.1191) | 344 (0.2251) | 534 (0.3495) | 679 (0.4444) |
| | **Union** | 176 (0.1152) | 369 (0.2415) | 533 (0.3488) | 675 (0.4418) | 170 (0.1113) | 353 (0.2310) | 535 (0.3501) | 692 (0.4529) |
| **EmbedPVP (Elboxembeddings)** | **GO** | 183 (0.1198) | 354 (0.2317) | 536 (0.3508) | 698 (0.4568) | 186 (0.1217) | 341 (0.2232) | 540 (0.3534) | 697 (0.4562) |
| | **HP** | 175 (0.1145) | 420 (0.2749) | 653 (0.4274) | 753 (0.4928) | 181 (0.1185) | 441 (0.2886) | 632 (0.4136) | 770 (0.5039) |
| | **MP** | 175 (0.1145) | 440 (0.2880) | 547 (0.3580) | 702 (0.4594) | 181 (0.1185) | 431 (0.2821) | 549 (0.3593) | 725 (0.4745) |
| | **UBERON** | 184 (0.1204) | 343 (0.2245) | 534 (0.3495) | 687 (0.4496) | 182 (0.1191) | 339 (0.2219) | 531 (0.3475) | 693 (0.4535) |
| | **Union** | 167 (0.1093) | 332 (0.2173) | 536 (0.3508) | 677 (0.4431) | 171 (0.1119) | 365 (0.2389) | 527 (0.3449) | 676 (0.4424) |
| **EmbedPVP (TransD)** | **GO** | 307 (0.2009) | 563 (0.3685) | 726 (0.4751) | 829 (0.5425) | 670 (0.4385) | 894 (0.5851) | 1006 (0.6584) | 1042 (0.6819) |
| | **HP** | **482 (0.3154)** | **846 (0.5537)** | **1007 (0.6590)** | **1056 (0.6911)** | 996 (0.6518) | 1230 (0.8050) | 1352 (0.8848) | **1391 (0.9103)** |
| | **MP** | 396 (0.2592) | 675 (0.4418) | 868 (0.5681) | 947 (0.6198) | 779 (0.5098) | 922 (0.6034) | 1031 (0.6747) | 1072 (0.7016) |
| | **UBERON** | 287 (0.1878) | 509 (0.3331) | 674 (0.4411) | 800 (0.5236) | 699 (0.4575) | 892 (0.5838) | 995 (0.6512) | 1023 (0.6695) |
| | **Union** | 409 (0.2677) | 639 (0.4182) | 833 (0.5452) | 928 (0.6073) | 899 (0.5884) | 1086 (0.7107) | 1158 (0.7579) | 1245 (0.8148) |
| **EmbedPVP (TransE)** | **GO** | 201 (0.1315) | 384 (0.2513) | 535 (0.3501) | 678 (0.4437) | 206 (0.1348) | 385 (0.2520) | 535 (0.3501) | 678 (0.4437) |
| | **HP** | 218 (0.1427) | 415 (0.2716) | 589 (0.3855) | 710 (0.4647) | 362 (0.2369) | 597 (0.3907) | 818 (0.5353) | 911 (0.5962) |
| | **MP** | 205 (0.1342) | 375 (0.2454) | 535 (0.3501) | 678 (0.4437) | 205 (0.1342) | 387 (0.2533) | 535 (0.3501) | 678 (0.4437) |
| | **UBERON** | 198 (0.1296) | 377 (0.2467) | 533 (0.3488) | 676 (0.4424) | 206 (0.1348) | 374 (0.2448) | 533 (0.3488) | 677 (0.4431) |
| | **Union** | 202 (0.1322) | 383 (0.2507) | 535 (0.3501) | 680 (0.4450) | 205 (0.1342) | 387 (0.2533) | 539 (0.3527) | 682 (0.4463) |
| **EmbedPVP (TransR)** | **GO** | 170 (0.1113) | 348 (0.2277) | 525 (0.3436) | 663 (0.4339) | 162 (0.1060) | 360 (0.2356) | 551 (0.3606) | 682 (0.4463) |
| | **HP** | 228 (0.1492) | 433 (0.2834) | 611 (0.3999) | 755 (0.4941) | 353 (0.2310) | 595 (0.3894) | 801 (0.5242) | 897 (0.5870) |
| | **MP** | 184 (0.1204) | 358 (0.2343) | 540 (0.3534) | 672 (0.4398) | 188 (0.1230) | 385 (0.2520) | 577 (0.3776) | 706 (0.4620) |
| | **UBERON** | 178 (0.1165) | 342 (0.2238) | 540 (0.3534) | 684 (0.4476) | 185 (0.1211) | 354 (0.2317) | 547 (0.3580) | 693 (0.4535) |
| | **Union** | 197 (0.1289) | 370 (0.2421) | 550 (0.3599) | 703 (0.4601) | 234 (0.1531) | 414 (0.2709) | 640 (0.4188) | 790 (0.5170) |
| **EmbedPVP (DL2vec)** | **GO** | 152 (0.0995) | 382 (0.2500) | 554 (0.3626) | 614 (0.4018) | 491 (0.3213) | 804 (0.5262) | 944 (0.6178) | 1010 (0.6610) |
| | **HP** | 362 (0.2369) | 666 (0.4359) | 787 (0.5151) | 826 (0.5406) | 1011 (0.6616) | 1300 (0.8508) | 1366 (0.8940) | 1384 (0.9058) |
| | **MP** | 255 (0.1669) | 491 (0.3213) | 639 (0.4182) | 701 (0.4588) | 639 (0.4182) | 914 (0.5982) | 1043 (0.6826) | 1106 (0.7238) |
| | **UBERON** | 174 (0.1139) | 390 (0.2552) | 498 (0.3259) | 556 (0.3639) | 539 (0.3527) | 801 (0.5242) | 904 (0.5916) | 940 (0.6152) |
| | **Union** | 358 (0.2343) | 636 (0.4162) | 771 (0.5046) | 824 (0.5393) | 950 (0.6217) | 1216 (0.7958) | 1310 (0.8573) | 1353 (0.8855) |
| **EmbedPVP (OWL2vec*)** | **GO** | 188 (0.1230) | 385 (0.2520) | 525 (0.3436) | 592 (0.3874) | 557 (0.3645) | 876 (0.5733) | 1011 (0.6616) | 1059 (0.6931) |
| | **HP** | 409 (0.2677) | 685 (0.4483) | 783 (0.5124) | 842 (0.5510) | 1026 (0.6715) | **1313 (0.8593)** | **1373 (0.8986)** | **1391 (0.9103)** |
| | **MP** | 222 (0.1453) | 470 (0.3076) | 618 (0.4045) | 677 (0.4431) | 665 (0.4352) | 965 (0.6315) | 1068 (0.6990) | 1116 (0.7304) |
| | **UBERON** | 158 (0.1034) | 379 (0.2480) | 474 (0.3102) | 525 (0.3436) | 577 (0.3776) | 800 (0.5236) | 888 (0.5812) | 937 (0.6132) |
| | **Union** | 375 (0.2454) | 650 (0.4254) | 787 (0.5151) | 835 (0.5465) | 959 (0.6276) | 1253 (0.8200) | 1325 (0.8671) | 1368 (0.8953) |

Table 1: EmbedPVP variant prediction results across several models and neuro-symbolic knowledge embeddings methods

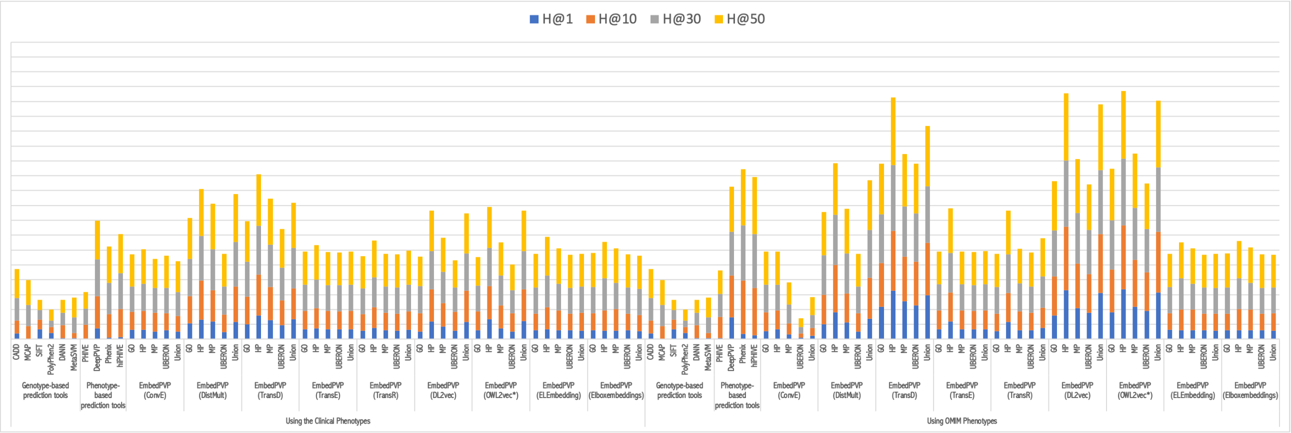


Figure 3: Recall at 1, 10, 30, and 50 using synthetic datasets with clinical phenotypes and OMIM phenotypes (PAVS dataset)

| | | **Exonic** | | | | **Non-Exonic** | | | |
|---|---|---|---|---|---|---|---|---|---|
| | | **H@1** | **H@10** | **H@30** | **H@50** | **H@1** | **H@10** | **H@30** | **H@50** |
| **Genotype-based prediction tools** | **CADD** | 116 (0.0995) | 205 (0.1758) | 397 (0.3405) | 521 (0.4468) | 8 (0.0270) | 78 (0.2635) | 99 (0.3345) | 107 (0.3615) |
| | **MCAP** | 4 (0.0034) | 261 (0.2238) | 442 (0.3791) | 511 (0.4383) | 0 (0.0) | 42 (0.1419) | 67 (0.2264) | 74 (0.25) |
| | **SIFT** | 201 (0.1724) | 201 (0.1724) | 201 (0.1724) | 201 (0.1724) | 12 (0.0405) | 12 (0.0405) | 12 (0.0405) | 12 (0.0405) |
| | **PolyPhen2** | 127 (0.1089) | 127 (0.1089) | 127 (0.1089) | 226 (0.1938) | 20 (0.0676) | 20 (0.0676) | 20 (0.0676) | 63 (0.2128) |
| | **DANN** | 21 (0.0180) | 263 (0.2256) | 263 (0.2256) | 263 (0.2256) | 0 (0.0) | 17 (0.0574) | 17 (0.0574) | 17 (0.0574) |
| | **MetaSVM** | 20 (0.0172) | 111 (0.0952) | 318 (0.2727) | 406 (0.3482) | 3 (0.0101) | 28 (0.0946) | 41 (0.1385) | 47 (0.1588) |
| **Phenotype-based prediction tools** | **PHIVE** | 169 (0.1449) | 304 (0.2607) | 331 (0.2839) | 347 (0.2976) | 32 (0.1081) | 46 (0.1554) | 49 (0.1655) | 49 (0.1655) |
| | **DeepPVP** | 201 (0.1724) | 600 (0.5146) | 689 (0.5909) | 720 (0.6175) | 9 (0.0304) | 34 (0.1149) | 43 (0.1453) | 44 (0.1486) |
| | **Phenix** | 419 (0.3593) | 564 (0.4837) | 675 (0.5789) | 709 (0.6081) | 79 (0.2669) | **117 (0.3953)** | **160 (0.5405)** | **175 (0.5912)** |
| | **hiPHIVE** | 383 (0.3285) | 581 (0.4983) | 674 (0.5780) | 709 (0.6081) | **80 (0.2703)** | 110 (0.3716) | 147 (0.4966) | 154 (0.5203) |
| **EmbedPVP (ConvE)** | **GO** | 194 (0.1664) | 317 (0.2719) | 455 (0.3902) | 590 (0.5060) | 4 (0.0135) | 55 (0.1858) | 74 (0.25) | 75 (0.2534) |
| | **HP** | 192 (0.1647) | 328 (0.2813) | 488 (0.4185) | 639 (0.5480) | 1 (0.0034) | 67 (0.2264) | 76 (0.2568) | 76 (0.2568) |
| | **MP** | 147 (0.1261) | 347 (0.2976) | 463 (0.3971) | 545 (0.4674) | 8 (0.0270) | 41 (0.1385) | 55 (0.1858) | 60 (0.2027) |
| | **UBERON** | 183 (0.1569) | 303 (0.2599) | 452 (0.3877) | 597 (0.5120) | 1 (0.0034) | 53 (0.1791) | 75 (0.2534) | 75 (0.2534) |
| | **Union** | 159 (0.1364) | 286 (0.2453) | 431 (0.3696) | 568 (0.4871) | 1 (0.0034) | 35 (0.1182) | 69 (0.2331) | 70 (0.2365) |
| **EmbedPVP (DistMult)** | **GO** | 289 (0.2479) | 505 (0.4331) | 691 (0.5926) | 774 (0.6638) | 36 (0.1216) | 64 (0.2162) | 78 (0.2635) | 83 (0.2804) |
| | **HP** | 365 (0.3130) | 734 (0.6295) | 834 (0.7153) | 881 (0.7556) | 37 (0.1250) | 85 (0.2872) | 101 (0.3412) | 109 (0.3682) |
| | **MP** | 333 (0.2856) | 587 (0.5034) | 772 (0.6621) | 862 (0.7393) | 35 (0.1182) | 64 (0.2162) | 80 (0.2703) | 87 (0.2939) |
| | **UBERON** | 140 (0.1201) | 312 (0.2676) | 507 (0.4348) | 615 (0.5274) | 10 (0.0338) | 53 (0.1791) | 68 (0.2297) | 73 (0.2466) |
| | **Union** | 330 (0.2830) | 664 (0.5695) | 838 (0.7187) | 904 (0.7753) | 27 (0.0912) | 70 (0.2365) | 87 (0.2939) | 96 (0.3243) |
| **EmbedPVP (ELEmbedding)** | **GO** | 183 (0.1569) | 305 (0.2616) | 470 (0.4031) | 618 (0.53) | 1 (0.0034) | 40 (0.1351) | 76 (0.2568) | 76 (0.2568) |
| | **HP** | 203 (0.1741) | 391 (0.3353) | 579 (0.4966) | 719 (0.6166) | 1 (0.0034) | 73 (0.2466) | 76 (0.2568) | 80 (0.2703) |
| | **MP** | 184 (0.1578) | 339 (0.2907) | 479 (0.4108) | 657 (0.5635) | 1 (0.0034) | 71 (0.2399) | 76 (0.2568) | 76 (0.2568) |
| | **UBERON** | 179 (0.1535) | 300 (0.2573) | 464 (0.3979) | 608 (0.5214) | 1 (0.0034) | 47 (0.1588) | 76 (0.2568) | 76 (0.2568) |
| | **Union** | 176 (0.1509) | 321 (0.2753) | 463 (0.3971) | 605 (0.5189) | 1 (0.0034) | 53 (0.1791) | 76 (0.2568) | 76 (0.2568) |
| **EmbedPVP (Elboxembeddings)** | **GO** | 183 (0.1569) | 311 (0.2667) | 466 (0.3997) | 628 (0.5386) | 1 (0.0034) | 48 (0.1622) | 76 (0.2568) | 76 (0.2568) |
| | **HP** | 175 (0.1501) | 357 (0.3062) | 583 (0.5) | 683 (0.5858) | 1 (0.0034) | 69 (0.2331) | 76 (0.2568) | 78 (0.2635) |
| | **MP** | 175 (0.1501) | 375 (0.3216) | 477 (0.4091) | 632 (0.5420) | 1 (0.0034) | 71 (0.2399) | 76 (0.2568) | 76 (0.2568) |
| | **UBERON** | 184 (0.1578) | 293 (0.2513) | 464 (0.3979) | 617 (0.5292) | 1 (0.0034) | 56 (0.1892) | 76 (0.2568) | 76 (0.2568) |
| | **Union** | 167 (0.1432) | 299 (0.2564) | 466 (0.3997) | 607 (0.5206) | 1 (0.0034) | 38 (0.1284) | 76 (0.2568) | 76 (0.2568) |
| **EmbedPVP (TransD)** | **GO** | 293 (0.2513) | 501 (0.4297) | 652 (0.5592) | 751 (0.6441) | 19 (0.0642) | 68 (0.2297) | 83 (0.2804) | 88 (0.2973) |
| | **HP** | **444 (0.3808)** | **759 (0.6509)** | **893 (0.7659)** | **930 (0.7976)** | 49 (0.1655) | 103 (0.3480) | 120 (0.4054) | 124 (0.4189) |
| | **MP** | 372 (0.3190) | 618 (0.5300) | 789 (0.6767) | 861 (0.7384) | 30 (0.1014) | 65 (0.2196) | 91 (0.3074) | 98 (0.3311) |
| | **UBERON** | 266 (0.2281) | 455 (0.3902) | 598 (0.5129) | 720 (0.6175) | 25 (0.0845) | 64 (0.2162) | 90 (0.3041) | 98 (0.3311) |
| | **Union** | 376 (0.3225) | 575 (0.4931) | 753 (0.6458) | 835 (0.7161) | 40 (0.1351) | 79 (0.2669) | 103 (0.3480) | 106 (0.3581) |
| **EmbedPVP (TransE)** | **GO** | 201 (0.1724) | 326 (0.2796) | 465 (0.3988) | 608 (0.5214) | 1 (0.0034) | 63 (0.2128) | 76 (0.2568) | 76 (0.2568) |
| | **HP** | 214 (0.1835) | 354 (0.3036) | 518 (0.4443) | 639 (0.5480) | 6 (0.0203) | 67 (0.2264) | 78 (0.2635) | 78 (0.2635) |
| | **MP** | 205 (0.1758) | 323 (0.2770) | 465 (0.3988) | 608 (0.5214) | 1 (0.0034) | 58 (0.1959) | 76 (0.2568) | 76 (0.2568) |
| | **UBERON** | 198 (0.1698) | 323 (0.2770) | 463 (0.3971) | 606 (0.5197) | 1 (0.0034) | 59 (0.1993) | 76 (0.2568) | 76 (0.2568) |
| | **Union** | 202 (0.1732) | 323 (0.2770) | 465 (0.3988) | 610 (0.5232) | 1 (0.0034) | 66 (0.2230) | 76 (0.2568) | 76 (0.2568) |
| **EmbedPVP (TransR)** | **GO** | 170 (0.1458) | 308 (0.2642) | 455 (0.3902) | 593 (0.5086) | 1 (0.0034) | 45 (0.1520) | 76 (0.2568) | 76 (0.2568) |
| | **HP** | 224 (0.1921) | 369 (0.3165) | 541 (0.4640) | 683 (0.5858) | 8 (0.0270) | 70 (0.2365) | 76 (0.2568) | 79 (0.2669) |
| | **MP** | 183 (0.1569) | 315 (0.2702) | 470 (0.4031) | 602 (0.5163) | 2 (0.0068) | 49 (0.1655) | 76 (0.2568) | 76 (0.2568) |
| | **UBERON** | 178 (0.1527) | 304 (0.2607) | 470 (0.4031) | 614 (0.5266) | 1 (0.0034) | 43 (0.1453) | 76 (0.2568) | 76 (0.2568) |
| | **Union** | 196 (0.1681) | 317 (0.2719) | 480 (0.4117) | 633 (0.5429) | 2 (0.0068) | 59 (0.1993) | 76 (0.2568) | 76 (0.2568) |
| **EmbedPVP (DL2vec)** | **GO** | 137 (0.1175) | 345 (0.2959) | 499 (0.4280) | 553 (0.4743) | 18 (0.0608) | 48 (0.1622) | 64 (0.2162) | 68 (0.2297) |
| | **HP** | 333 (0.2856) | 571 (0.4897) | 676 (0.5798) | 707 (0.6063) | 40 (0.1351) | 87 (0.2939) | 96 (0.3243) | 101 (0.3412) |
| | **MP** | 234 (0.2007) | 443 (0.3799) | 572 (0.4906) | 628 (0.5386) | 26 (0.0878) | 58 (0.1959) | 92 (0.3108) | 101 (0.3412) |
| | **UBERON** | 161 (0.1381) | 349 (0.2993) | 439 (0.3765) | 490 (0.4202) | 19 (0.0642) | 50 (0.1689) | 63 (0.2128) | 66 (0.2230) |
| | **Union** | 325 (0.2787) | 557 (0.4777) | 671 (0.5755) | 714 (0.6123) | 42 (0.1419) | 79 (0.2669) | 94 (0.3176) | 104 (0.3514) |
| **EmbedPVP (OWL2vec*)** | **GO** | 171 (0.1467) | 357 (0.3062) | 476 (0.4082) | 535 (0.4588) | 23 (0.0777) | 39 (0.1318) | 56 (0.1892) | 65 (0.2196) |
| | **HP** | 375 (0.3216) | 587 (0.5034) | 673 (0.5772) | 721 (0.6184) | 50 (0.1689) | 88 (0.2973) | 101 (0.3412) | 107 (0.3615) |
| | **MP** | 206 (0.1767) | 419 (0.3593) | 549 (0.4708) | 600 (0.5146) | 22 (0.0743) | 54 (0.1824) | 75 (0.2534) | 93 (0.3142) |
| | **UBERON** | 147 (0.1261) | 348 (0.2985) | 434 (0.3722) | 478 (0.4099) | 16 (0.0541) | 40 (0.1351) | 50 (0.1689) | 54 (0.1824) |
| | **Union** | 346 (0.2967) | 579 (0.4966) | 683 (0.5858) | 728 (0.6244) | 41 (0.1385) | 72 (0.2432) | 88 (0.2973) | 89 (0.3007) |

Table 2: EmbedPVP variant prediction results across several models for the exonic and non-exonic variants

##### 2.1.2 Phenopackets dataset

Table 3, Figure 4

| | | Using the Clinical Phenotypes | | | | Using OMIM Phenotypes | | | |
|---|---|---|---|---|---|---|---|---|---|
| | | **H@1** | **H@10** | **H@30** | **H@50** | **H@1** | **H@10** | **H@30** | **H@50** |
| **Genotype-based prediction tools** | **CADD** | 51 (0.1328) | 104 (0.2708) | 152 (0.3958) | 184 (0.4792) | 51 (0.1328) | 104 (0.2708) | 152 (0.3958) | 184 (0.4792) |
| | **MCAP** | 2 (0.0052) | 68 (0.1771) | 105 (0.2734) | 122 (0.3177) | 2 (0.0052) | 68 (0.1771) | 105 (0.2734) | 122 (0.3177) |
| | **SIFT** | 69 (0.1797) | 69 (0.1797) | 69 (0.1797) | 69 (0.1797) | 69 (0.1797) | 69 (0.1797) | 69 (0.1797) | 69 (0.1797) |
| | **PolyPhen2** | 51 (0.1328) | 51 (0.1328) | 51 (0.1328) | 71 (0.1849) | 51 (0.1328) | 51 (0.1328) | 51 (0.1328) | 71 (0.1849) |
| | **DANN** | 4 (0.0104) | 81 (0.2109) | 81 (0.2109) | 81 (0.2109) | 4 (0.0104) | 81 (0.2109) | 81 (0.2109) | 81 (0.2109) |
| | **MetaSVM** | 12 (0.0312) | 37 (0.0964) | 80 (0.2083) | 101 (0.2630) | 12 (0.0312) | 37 (0.0964) | 80 (0.2083) | 101 (0.2630) |
| **Phenotype-based prediction tools** | **PHIVE** | 48 (0.1250) | 80 (0.2083) | 87 (0.2266) | 87 (0.2266) | 53 (0.1380) | 94 (0.2448) | 100 (0.2604) | 100 (0.2604) |
| | **DeepPVP** | 135 (0.3516) | 204 (0.5312) | 220 (0.5729) | 228 (0.5938) | 193 (0.5026) | 232 (0.6042) | 233 (0.6068) | 244 (0.6354) |
| | **Phenix** | **202 (0.5260)** | **250 (0.6510)** | **293 (0.7630)** | **308 (0.8021)** | **359 (0.9349)** | **372 (0.9688)** | **378 (0.9844)** | **378 (0.9844)** |
| | **hiPHIVE** | 193 (0.5026) | 245 (0.6380) | 264 (0.6875) | 278 (0.7240) | 285 (0.7422) | 342 (0.8906) | 371 (0.9661) | 377 (0.9818) |
| **EmbedPVP (ConvE)** | **GO** | 6 (0.0156) | 18 (0.0469) | 39 (0.1016) | 47 (0.1224) | 17 (0.0443) | 50 (0.1302) | 84 (0.2188) | 105 (0.2734) |
| | **HP** | 38 (0.0990) | 93 (0.2422) | 141 (0.3672) | 172 (0.4479) | 40 (0.1042) | 104 (0.2708) | 153 (0.3984) | 183 (0.4766) |
| | **MP** | 17 (0.0443) | 62 (0.1615) | 101 (0.2630) | 122 (0.3177) | 6 (0.0156) | 34 (0.0885) | 56 (0.1458) | 68 (0.1771) |
| | **UBERON** | 43 (0.1120) | 92 (0.2396) | 139 (0.3620) | 176 (0.4583) | 46 (0.1198) | 96 (0.25) | 152 (0.3958) | 185 (0.4818) |
| | **Union** | 46 (0.1198) | 96 (0.25) | 148 (0.3854) | 182 (0.4740) | 16 (0.0417) | 40 (0.1042) | 80 (0.2083) | 104 (0.2708) |
| **EmbedPVP (DistMult)** | **GO** | 60 (0.1562) | 150 (0.3906) | 220 (0.5729) | 242 (0.6302) | 89 (0.2318) | 145 (0.3776) | 200 (0.5208) | 221 (0.5755) |
| | **HP** | 102 (0.2656) | 197 (0.5130) | 218 (0.5677) | 221 (0.5755) | 112 (0.2917) | 210 (0.5469) | 223 (0.5807) | 225 (0.5859) |
| | **MP** | 76 (0.1979) | 147 (0.3828) | 203 (0.5286) | 229 (0.5964) | 62 (0.1615) | 142 (0.3698) | 200 (0.5208) | 236 (0.6146) |
| | **UBERON** | 42 (0.1094) | 107 (0.2786) | 151 (0.3932) | 189 (0.4922) | 41 (0.1068) | 108 (0.2812) | 167 (0.4349) | 208 (0.5417) |
| | **Union** | 38 (0.0990) | 152 (0.3958) | 202 (0.5260) | 216 (0.5625) | 75 (0.1953) | 158 (0.4115) | 207 (0.5391) | 219 (0.5703) |
| **EmbedPVP (ELEmbedding)** | **GO** | 47 (0.1224) | 93 (0.2422) | 155 (0.4036) | 183 (0.4766) | 46 (0.1198) | 95 (0.2474) | 154 (0.4010) | 187 (0.4870) |
| | **HP** | 43 (0.1120) | 119 (0.3099) | 169 (0.4401) | 198 (0.5156) | 40 (0.1042) | 109 (0.2839) | 168 (0.4375) | 192 (0.5000) |
| | **MP** | 45 (0.1172) | 108 (0.2812) | 154 (0.4010) | 194 (0.5052) | 46 (0.1198) | 106 (0.2760) | 153 (0.3984) | 190 (0.4948) |
| | **UBERON** | 47 (0.1224) | 92 (0.2396) | 151 (0.3932) | 186 (0.4844) | 46 (0.1198) | 88 (0.2292) | 153 (0.3984) | 186 (0.4844) |
| | **Union** | 46 (0.1198) | 93 (0.2422) | 148 (0.3854) | 183 (0.4766) | 46 (0.1198) | 94 (0.2448) | 151 (0.3932) | 181 (0.4714) |
| **EmbedPVP (Elboxembeddings)** | **GO** | 47 (0.1224) | 97 (0.2526) | 151 (0.3932) | 186 (0.4844) | 47 (0.1224) | 96 (0.25) | 145 (0.3776) | 181 (0.4714) |
| | **HP** | 42 (0.1094) | 114 (0.2969) | 168 (0.4375) | 194 (0.5052) | 40 (0.1042) | 115 (0.2995) | 172 (0.4479) | 198 (0.5156) |
| | **MP** | 46 (0.1198) | 105 (0.2734) | 157 (0.4089) | 189 (0.4922) | 41 (0.1068) | 112 (0.2917) | 159 (0.4141) | 187 (0.4870) |
| | **UBERON** | 45 (0.1172) | 99 (0.2578) | 150 (0.3906) | 182 (0.4740) | 46 (0.1198) | 89 (0.2318) | 153 (0.3984) | 180 (0.4688) |
| | **Union** | 46 (0.1198) | 94 (0.2448) | 151 (0.3932) | 184 (0.4792) | 46 (0.1198) | 99 (0.2578) | 142 (0.3698) | 179 (0.4661) |
| **EmbedPVP (TransD)** | **GO** | 101 (0.2630) | 177 (0.4609) | 228 (0.5938) | 250 (0.6510) | 140 (0.3646) | 207 (0.5391) | 237 (0.6172) | 257 (0.6693) |
| | **HP** | 160 (0.4167) | 217 (0.5651) | 230 (0.5990) | 240 (0.6250) | 212 (0.5521) | 244 (0.6354) | 267 (0.6953) | 272 (0.7083) |
| | **MP** | 134 (0.3490) | 194 (0.5052) | 235 (0.6120) | 252 (0.6562) | 175 (0.4557) | 215 (0.5599) | 233 (0.6068) | 248 (0.6458) |
| | **UBERON** | 120 (0.3125) | 202 (0.5260) | 237 (0.6172) | 253 (0.6589) | 156 (0.4062) | 220 (0.5729) | 240 (0.6250) | 258 (0.6719) |
| | **Union** | 136 (0.3542) | 217 (0.5651) | 247 (0.6432) | 257 (0.6693) | 199 (0.5182) | 230 (0.5990) | 256 (0.6667) | 265 (0.6901) |
| **EmbedPVP (TransE)** | **GO** | 49 (0.1276) | 102 (0.2656) | 152 (0.3958) | 184 (0.4792) | 50 (0.1302) | 101 (0.2630) | 152 (0.3958) | 184 (0.4792) |
| | **HP** | 51 (0.1328) | 105 (0.2734) | 160 (0.4167) | 191 (0.4974) | 97 (0.2526) | 163 (0.4245) | 220 (0.5729) | 242 (0.6302) |
| | **MP** | 51 (0.1328) | 102 (0.2656) | 152 (0.3958) | 184 (0.4792) | 50 (0.1302) | 101 (0.2630) | 152 (0.3958) | 184 (0.4792) |
| | **UBERON** | 49 (0.1276) | 100 (0.2604) | 152 (0.3958) | 184 (0.4792) | 50 (0.1302) | 99 (0.2578) | 152 (0.3958) | 184 (0.4792) |
| | **Union** | 50 (0.1302) | 99 (0.2578) | 152 (0.3958) | 184 (0.4792) | 50 (0.1302) | 101 (0.2630) | 154 (0.4010) | 185 (0.4818) |
| **EmbedPVP (TransR)** | **GO** | 40 (0.1042) | 94 (0.2448) | 146 (0.3802) | 184 (0.4792) | 34 (0.0885) | 93 (0.2422) | 145 (0.3776) | 185 (0.4818) |
| | **HP** | 57 (0.1484) | 121 (0.3151) | 183 (0.4766) | 210 (0.5469) | 103 (0.2682) | 165 (0.4297) | 215 (0.5599) | 237 (0.6172) |
| | **MP** | 45 (0.1172) | 99 (0.2578) | 152 (0.3958) | 185 (0.4818) | 47 (0.1224) | 100 (0.2604) | 158 (0.4115) | 190 (0.4948) |
| | **UBERON** | 45 (0.1172) | 95 (0.2474) | 159 (0.4141) | 184 (0.4792) | 46 (0.1198) | 98 (0.2552) | 156 (0.4062) | 190 (0.4948) |
| | **Union** | 53 (0.1380) | 109 (0.2839) | 161 (0.4193) | 199 (0.5182) | 60 (0.1562) | 120 (0.3125) | 178 (0.4635) | 211 (0.5495) |
| **EmbedPVP (DL2vec)** | **GO** | 71 (0.1849) | 142 (0.3698) | 179 (0.4661) | 198 (0.5156) | 139 (0.3620) | 204 (0.5312) | 233 (0.6068) | 241 (0.6276) |
| | **HP** | 161 (0.4193) | 216 (0.5625) | 233 (0.6068) | 237 (0.6172) | 233 (0.6068) | 270 (0.7031) | 274 (0.7135) | 276 (0.7188) |
| | **MP** | 86 (0.2240) | 160 (0.4167) | 183 (0.4766) | 200 (0.5208) | 169 (0.4401) | 207 (0.5391) | 224 (0.5833) | 246 (0.6406) |
| | **UBERON** | 83 (0.2161) | 150 (0.3906) | 178 (0.4635) | 193 (0.5026) | 163 (0.4245) | 207 (0.5391) | 223 (0.5807) | 230 (0.5990) |
| | **Union** | 122 (0.3177) | 193 (0.5026) | 219 (0.5703) | 235 (0.6120) | 208 (0.5417) | 252 (0.6562) | 269 (0.7005) | 275 (0.7161) |
| **EmbedPVP (OWL2vec*)** | **GO** | 70 (0.1823) | 142 (0.3698) | 181 (0.4714) | 204 (0.5312) | 150 (0.3906) | 211 (0.5495) | 234 (0.6094) | 252 (0.6562) |
| | **HP** | 156 (0.4062) | 213 (0.5547) | 226 (0.5885) | 236 (0.6146) | 235 (0.6120) | 270 (0.7031) | 276 (0.7188) | 277 (0.7214) |
| | **MP** | 88 (0.2292) | 164 (0.4271) | 190 (0.4948) | 200 (0.5208) | 165 (0.4297) | 205 (0.5339) | 220 (0.5729) | 230 (0.5990) |
| | **UBERON** | 75 (0.1953) | 144 (0.3750) | 173 (0.4505) | 187 (0.4870) | 155 (0.4036) | 202 (0.5260) | 220 (0.5729) | 231 (0.6016) |
| | **Union** | 119 (0.3099) | 197 (0.5130) | 211 (0.5495) | 220 (0.5729) | 220 (0.5729) | 264 (0.6875) | 281 (0.7318) | 285 (0.7422) |

Table 3: EmbedPVP variant prediction results across several models using Phenopackets dataset

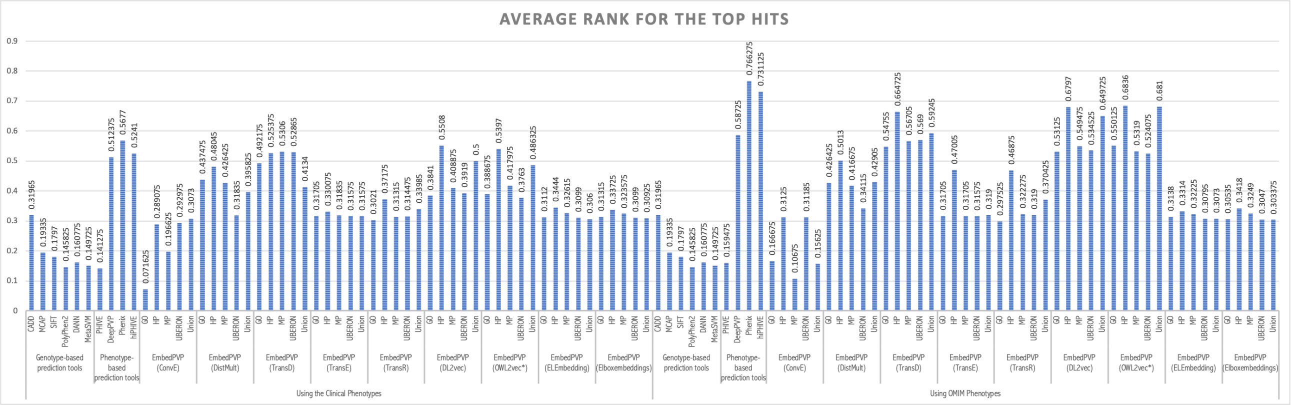


Figure 4: Average ranks for recall at 1, 10, 30, and 50 using synthetic datasets with clinical phenotypes and OMIM phenotypes (Phenopackets)

##### 2.1.3 ClinVar time-split

The results are in Table 4. Tables 5 and 6 show the results of using novel and known genes or diseases. Table 7 provides the evaluation for exonic and non-exonic variants.

| | | H@1 | H@10 | H@30 | H@50 |
|---|---|---|---|---|---|
| **Genotype-based prediction tools** | **CADD** | 239 (0.2209) | 505 (0.4667) | 600 (0.5545) | 688 (0.6359) |
| | **MCAP** | 11 (0.0102) | 125 (0.1155) | 201 (0.1858) | 244 (0.2255) |
| | **SIFT** | 198 (0.1830) | 198 (0.1830) | 198 (0.1830) | 198 (0.1830) |
| | **PolyPhen2** | 112 (0.1035) | 112 (0.1035) | 112 (0.1035) | 182 (0.1682) |
| | **DANN** | 7 (0.0065) | 172 (0.1590) | 172 (0.1590) | 172 (0.1590) |
| | **MetaSVM** | 20 (0.0185) | 74 (0.0684) | 139 (0.1285) | 177 (0.1636) |
| **Phenotype-based prediction tools** | **PHIVE** | 175 (0.1617) | 284 (0.2625) | 298 (0.2754) | 301 (0.2782) |
| | **deeppvp** | 396 (0.3660) | 532 (0.4917) | 560 (0.5176) | 572 (0.5287) |
| | **Phenix** | 118 (0.1091) | 567 (0.5240) | 620 (0.5730) | 633 (0.5850) |
| | **hiPHIVE** | 487 (0.4501) | 586 (0.5416) | 641 (0.5924) | 662 (0.6118) |
| **EmbedPVP (ConvE)** | **GO** | 102 (0.0943) | 272 (0.2514) | 385 (0.3558) | 435 (0.4020) |
| | **HP** | 220 (0.2033) | 501 (0.4630) | 597 (0.5518) | 686 (0.6340) |
| | **MP** | 208 (0.1922) | 460 (0.4251) | 579 (0.5351) | 662 (0.6118) |
| | **UBERON** | 205 (0.1895) | 504 (0.4658) | 599 (0.5536) | 688 (0.6359) |
| | **Union** | 188 (0.1738) | 401 (0.3706) | 564 (0.5213) | 662 (0.6118) |
| **EmbedPVP (DistMult)** | **GO** | 314 (0.2902) | 480 (0.4436) | 628 (0.5804) | 694 (0.6414) |
| | **HP** | 345 (0.3189) | 571 (0.5277) | 611 (0.5647) | 623 (0.5758) |
| | **MP** | 350 (0.3235) | 489 (0.4519) | 663 (0.6128) | 744 (0.6876) |
| | **UBERON** | 214 (0.1978) | 461 (0.4261) | 608 (0.5619) | 689 (0.6368) |
| | **Union** | 246 (0.2274) | 498 (0.4603) | 616 (0.5693) | 661 (0.6109) |
| **EmbedPVP (ELEmbedding)** | **GO** | 215 (0.1987) | 454 (0.4196) | 600 (0.5545) | 695 (0.6423) |
| | **HP** | 189 (0.1747) | 515 (0.4760) | 616 (0.5693) | 694 (0.6414) |
| | **MP** | 209 (0.1932) | 518 (0.4787) | 597 (0.5518) | 706 (0.6525) |
| | **UBERON** | 205 (0.1895) | 466 (0.4307) | 598 (0.5527) | 691 (0.6386) |
| | **Union** | 206 (0.1904) | 454 (0.4196) | 595 (0.5499) | 691 (0.6386) |
| **EmbedPVP (Elboxembeddings)** | **GO** | 202 (0.1867) | 464 (0.4288) | 599 (0.5536) | 693 (0.6405) |
| | **HP** | 180 (0.1664) | 505 (0.4667) | 614 (0.5675) | 687 (0.6349) |
| | **MP** | 208 (0.1922) | 500 (0.4621) | 602 (0.5564) | 703 (0.6497) |
| | **UBERON** | 213 (0.1969) | 442 (0.4085) | 590 (0.5453) | 685 (0.6331) |
| | **Union** | 204 (0.1885) | 440 (0.4067) | 599 (0.5536) | 684 (0.6322) |
| **EmbedPVP (TransD)** | **GO** | 463 (0.4279) | 646 (0.5970) | **798 (0.7375)** | **846 (0.7819)** |
| | **HP** | **554 (0.5120)** | 633 (0.5850) | 647 (0.5980) | 660 (0.6100) |
| | **MP** | 526 (0.4861) | 681 (0.6294) | 777 (0.7181) | 815 (0.7532) |
| | **UBERON** | 485 (0.4482) | 633 (0.5850) | 748 (0.6913) | 808 (0.7468) |
| | **Union** | 537 (0.4963) | **691 (0.6386)** | 768 (0.7098) | 808 (0.7468) |
| **EmbedPVP (TransE)** | **GO** | 232 (0.2144) | 489 (0.4519) | 600 (0.5545) | 688 (0.6359) |
| | **HP** | 374 (0.3457) | 593 (0.5481) | 729 (0.6738) | 820 (0.7579) |
| | **MP** | 232 (0.2144) | 486 (0.4492) | 600 (0.5545) | 688 (0.6359) |
| | **UBERON** | 226 (0.2089) | 484 (0.4473) | 596 (0.5508) | 688 (0.6359) |
| | **Union** | 231 (0.2135) | 496 (0.4584) | 601 (0.5555) | 689 (0.6368) |
| **EmbedPVP (TransR)** | **GO** | 204 (0.1885) | 453 (0.4187) | 603 (0.5573) | 692 (0.6396) |
| | **HP** | 369 (0.3410) | 578 (0.5342) | 709 (0.6553) | 812 (0.7505) |
| | **MP** | 200 (0.1848) | 474 (0.4381) | 607 (0.5610) | 700 (0.6470) |
| | **UBERON** | 221 (0.2043) | 468 (0.4325) | 604 (0.5582) | 701 (0.6479) |
| | **Union** | 264 (0.2440) | 513 (0.4741) | 623 (0.5758) | 744 (0.6876) |
| **EmbedPVP (DL2vec)** | **GO** | 414 (0.3826) | 554 (0.5120) | 623 (0.5758) | 658 (0.6081) |
| | **HP** | 542 (0.5009) | 597 (0.5518) | 607 (0.5610) | 613 (0.5665) |
| | **MP** | 441 (0.4076) | 576 (0.5323) | 632 (0.5841) | 656 (0.6063) |
| | **UBERON** | 413 (0.3817) | 574 (0.5305) | 642 (0.5933) | 675 (0.6238) |
| | **Union** | 511 (0.4723) | 605 (0.5591) | 638 (0.5896) | 669 (0.6183) |
| **EmbedPVP (OWL2vec*)** | **GO** | 456 (0.4214) | 609 (0.5628) | 680 (0.6285) | 703 (0.6497) |
| | **HP** | 547 (0.5055) | 595 (0.5499) | 612 (0.5656) | 625 (0.5776) |
| | **MP** | 483 (0.4464) | 600 (0.5545) | 660 (0.6100) | 691 (0.6386) |
| | **UBERON** | 426 (0.3937) | 576 (0.5323) | 624 (0.5767) | 655 (0.6054) |
| | **Union** | 527 (0.4871) | 610 (0.5638) | 661 (0.6109) | 679 (0.6275) |

Table 4: EmbedPVP variant prediction results across several models using ClinVar dataset.

| | | Novel Genes and Diseases | | | | Novel Genes and Known Diseases | | | |
|---|---|---|---|---|---|---|---|---|---|
| | | **H@1** | **H@10** | **H@30** | **H@50** | **H@1** | **H@10** | **H@30** | **H@50** |
| **Genotype-based prediction tools** | **CADD** | 68 (0.1498) | 155 (0.3414) | 206 (0.4537) | 263 (0.5793) | 8 (0.2581) | 14 (0.4516) | 14 (0.4516) | 16 (0.5161) |
| | **MCAP** | 1 (0.0022) | 33 (0.0727) | 77 (0.1696) | 102 (0.2247) | 2 (0.0645) | 5 (0.1613) | 6 (0.1935) | 7 (0.2258) |
| | **SIFT** | 102 (0.2247) | 102 (0.2247) | 102 (0.2247) | 102 (0.2247) | 7 (0.2258) | 7 (0.2258) | 7 (0.2258) | 7 (0.2258) |
| | **PolyPhen2** | 59 (0.13) | 59 (0.13) | 59 (0.13) | 100 (0.2203) | 4 (0.1290) | 4 (0.1290) | 4 (0.1290) | 5 (0.1613) |
| | **DANN** | 5 (0.0110) | 108 (0.2379) | 108 (0.2379) | 108 (0.2379) | 0 (0.0) | 6 (0.1935) | 6 (0.1935) | 6 (0.1935) |
| | **MetaSVM** | 4 (0.0088) | 28 (0.0617) | 43 (0.0947) | 59 (0.1300) | 0 (0.0) | 3 (0.0968) | 6 (0.1935) | 7 (0.2258) |
| **Phenotype-based prediction tools** | **PHIVE** | 32 (0.0705) | 55 (0.1211) | 55 (0.1211) | 55 (0.1211) | 3 (0.0968) | 4 (0.1290) | 4 (0.1290) | 4 (0.1290) |
| | **DeepPVP** | 37 (0.0815) | 59 (0.1300) | 70 (0.1542) | 77 (0.1696) | **20 (0.6452)** | **23 (0.7419)** | **24 (0.7742)** | **24 (0.7742)** |
| | **Phenix** | 15 (0.0330) | 78 (0.1718) | 82 (0.1806) | 89 (0.1960) | 4 (0.1290) | 21 (0.6774) | 22 (0.7097) | 22 (0.7097) |
| | **hiPHIVE** | 77 (0.1696) | 100 (0.2203) | 119 (0.2621) | 121 (0.2665) | 13 (0.4194) | 18 (0.5806) | 19 (0.6129) | 22 (0.7097) |
| **EmbedPVP (ConvE)** | **GO** | 28 (0.0617) | 83 (0.1828) | 125 (0.2753) | 152 (0.3348) | 3 (0.0968) | 5 (0.1613) | 5 (0.1613) | 5 (0.1613) |
| | **HP** | 57 (0.1256) | 154 (0.3392) | 205 (0.4515) | 261 (0.5749) | 7 (0.2258) | 14 (0.4516) | 14 (0.4516) | 16 (0.5161) |
| | **MP** | 52 (0.1145) | 133 (0.2930) | 195 (0.4295) | 241 (0.5308) | 7 (0.2258) | 13 (0.4194) | 14 (0.4516) | 15 (0.4839) |
| | **UBERON** | 58 (0.1278) | 154 (0.3392) | 205 (0.4515) | 263 (0.5793) | 7 (0.2258) | 14 (0.4516) | 14 (0.4516) | 16 (0.5161) |
| | **Union** | 51 (0.1123) | 116 (0.2555) | 190 (0.4185) | 243 (0.5352) | 5 (0.1613) | 9 (0.2903) | 13 (0.4194) | 15 (0.4839) |
| **EmbedPVP (DistMult)** | **GO** | 33 (0.0727) | 77 (0.1696) | 143 (0.3150) | 173 (0.3811) | 4 (0.1290) | 11 (0.3548) | 14 (0.4516) | 15 (0.4839) |
| | **HP** | 25 (0.0551) | 46 (0.1013) | 51 (0.1123) | 52 (0.1145) | 13 (0.4194) | 20 (0.6452) | 21 (0.6774) | 21 (0.6774) |
| | **MP** | 53 (0.1167) | 106 (0.2335) | 186 (0.4097) | 224 (0.4934) | 8 (0.2581) | 10 (0.3226) | 14 (0.4516) | 17 (0.5484) |
| | **UBERON** | 32 (0.0705) | 121 (0.2665) | 193 (0.4251) | 226 (0.4978) | 5 (0.1613) | 10 (0.3226) | 12 (0.3871) | 12 (0.3871) |
| | **Union** | 15 (0.0330) | 55 (0.1211) | 91 (0.2004) | 111 (0.2445) | 7 (0.2258) | 14 (0.4516) | 19 (0.6129) | 20 (0.6452) |
| **EmbedPVP (ELEmbedding)** | **GO** | 60 (0.1322) | 138 (0.3040) | 208 (0.4581) | 268 (0.5903) | 8 (0.2581) | 11 (0.3548) | 14 (0.4516) | 16 (0.5161) |
| | **HP** | 30 (0.0661) | 141 (0.3106) | 201 (0.4427) | 231 (0.5088) | 7 (0.2258) | 13 (0.4194) | 16 (0.5161) | 17 (0.5484) |
| | **MP** | 62 (0.1366) | 159 (0.3502) | 204 (0.4493) | 264 (0.5815) | 7 (0.2258) | 14 (0.4516) | 14 (0.4516) | 16 (0.5161) |
| | **UBERON** | 56 (0.1233) | 143 (0.3150) | 206 (0.4537) | 264 (0.5815) | 7 (0.2258) | 11 (0.3548) | 14 (0.4516) | 16 (0.5161) |
| | **Union** | 57 (0.1256) | 141 (0.3106) | 203 (0.4471) | 266 (0.5859) | 7 (0.2258) | 11 (0.3548) | 14 (0.4516) | 16 (0.5161) |
| **EmbedPVP (Elboxembeddings)** | **GO** | 56 (0.1233) | 140 (0.3084) | 207 (0.4559) | 265 (0.5837) | 7 (0.2258) | 14 (0.4516) | 14 (0.4516) | 16 (0.5161) |
| | **HP** | 30 (0.0661) | 141 (0.3106) | 201 (0.4427) | 232 (0.5110) | 7 (0.2258) | 13 (0.4194) | 15 (0.4839) | 16 (0.5161) |
| | **MP** | 55 (0.1211) | 150 (0.3304) | 210 (0.4626) | 262 (0.5771) | 7 (0.2258) | 14 (0.4516) | 14 (0.4516) | 16 (0.5161) |
| | **UBERON** | 61 (0.1344) | 132 (0.2907) | 199 (0.4383) | 261 (0.5749) | 7 (0.2258) | 12 (0.3871) | 14 (0.4516) | 16 (0.5161) |
| | **Union** | 57 (0.1256) | 134 (0.2952) | 208 (0.4581) | 262 (0.5771) | 7 (0.2258) | 13 (0.4194) | 14 (0.4516) | 16 (0.5161) |
| **EmbedPVP (TransD)** | **GO** | 51 (0.1123) | 138 (0.3040) | 234 (0.5154) | 272 (0.5991) | 10 (0.3226) | 11 (0.3548) | 17 (0.5484) | 21 (0.6774) |
| | **HP** | 43 (0.0947) | 47 (0.1035) | 50 (0.1101) | 59 (0.13) | **20 (0.6452)** | 21 (0.6774) | 21 (0.6774) | 21 (0.6774) |
| | **MP** | 50 (0.1101) | 127 (0.2797) | 199 (0.4383) | 229 (0.5044) | 6 (0.1935) | 12 (0.3871) | 14 (0.4516) | 16 (0.5161) |
| | **UBERON** | 27 (0.0595) | 95 (0.2093) | 175 (0.3855) | 223 (0.4912) | 2 (0.0645) | 8 (0.2581) | 11 (0.3548) | 14 (0.4516) |
| | **Union** | 51 (0.1123) | 119 (0.2621) | 180 (0.3965) | 213 (0.4692) | **20 (0.6452)** | 22 (0.7097) | 23 (0.7419) | **24 (0.7742)** |
| **EmbedPVP (TransE)** | **GO** | 61 (0.1344) | 142 (0.3128) | 206 (0.4537) | 263 (0.5793) | 8 (0.2581) | 13 (0.4194) | 14 (0.4516) | 16 (0.5161) |
| | **HP** | **78 (0.1718)** | **164 (0.3612)** | **222 (0.4890)** | **278 (0.6123)** | 12 (0.3871) | 18 (0.5806) | 22 (0.7097) | 22 (0.7097) |
| | **MP** | 62 (0.1366) | 138 (0.3040) | 206 (0.4537) | 263 (0.5793) | 7 (0.2258) | 13 (0.4194) | 14 (0.4516) | 16 (0.5161) |
| | **UBERON** | 57 (0.1256) | 137 (0.3018) | 203 (0.4471) | 263 (0.5793) | 7 (0.2258) | 13 (0.4194) | 14 (0.4516) | 16 (0.5161) |
| | **Union** | 61 (0.1344) | 146 (0.3216) | 206 (0.4537) | 263 (0.5793) | 8 (0.2581) | 14 (0.4516) | 14 (0.4516) | 16 (0.5161) |
| **EmbedPVP (TransR)** | **GO** | 65 (0.1432) | 151 (0.3326) | 213 (0.4692) | 267 (0.5881) | 7 (0.2258) | 11 (0.3548) | 14 (0.4516) | 16 (0.5161) |
| | **HP** | 77 (0.1696) | 158 (0.3480) | 216 (0.4758) | 277 (0.6101) | 16 (0.5161) | 20 (0.6452) | 22 (0.7097) | 22 (0.7097) |
| | **MP** | 52 (0.1145) | 143 (0.3150) | 210 (0.4626) | 267 (0.5881) | 7 (0.2258) | 13 (0.4194) | 14 (0.4516) | 17 (0.5484) |
| | **UBERON** | 60 (0.1322) | 142 (0.3128) | 207 (0.4559) | 265 (0.5837) | 7 (0.2258) | 12 (0.3871) | 14 (0.4516) | 16 (0.5161) |
| | **Union** | 68 (0.1498) | 150 (0.3304) | 213 (0.4692) | 271 (0.5969) | 10 (0.3226) | 13 (0.4194) | 15 (0.4839) | 19 (0.6129) |
| **EmbedPVP (DL2vec)** | **GO** | 34 (0.0749) | 60 (0.1322) | 88 (0.1938) | 109 (0.2401) | 18 (0.5806) | 19 (0.6129) | 20 (0.6452) | 21 (0.6774) |
| | **HP** | 44 (0.0969) | 46 (0.1013) | 46 (0.1013) | 46 (0.1013) | **20 (0.6452)** | 21 (0.6774) | 21 (0.6774) | 21 (0.6774) |
| | **MP** | 37 (0.0815) | 54 (0.1189) | 76 (0.1674) | 89 (0.1960) | 13 (0.4194) | 19 (0.6129) | 20 (0.6452) | 21 (0.6774) |
| | **UBERON** | 25 (0.0551) | 55 (0.1211) | 90 (0.1982) | 110 (0.2423) | 14 (0.4516) | 17 (0.5484) | 20 (0.6452) | 20 (0.6452) |
| | **Union** | 44 (0.0969) | 62 (0.1366) | 76 (0.1674) | 96 (0.2115) | 19 (0.6129) | 20 (0.6452) | 21 (0.6774) | 21 (0.6774) |
| **EmbedPVP (OWL2vec*)** | **GO** | 40 (0.0881) | 82 (0.1806) | 123 (0.2709) | 136 (0.2996) | 17 (0.5484) | 21 (0.6774) | 23 (0.7419) | 23 (0.7419) |
| | **HP** | 45 (0.0991) | 48 (0.1057) | 48 (0.1057) | 48 (0.1057) | **20 (0.6452)** | 21 (0.6774) | 21 (0.6774) | 21 (0.6774) |
| | **MP** | 42 (0.0925) | 68 (0.1498) | 93 (0.2048) | 113 (0.2489) | 17 (0.5484) | 20 (0.6452) | 21 (0.6774) | 21 (0.6774) |
| | **UBERON** | 30 (0.0661) | 54 (0.1189) | 70 (0.1542) | 82 (0.1806) | 13 (0.4194) | 20 (0.6452) | 21 (0.6774) | 22 (0.7097) |
| | **Union** | 46 (0.1013) | 57 (0.1256) | 76 (0.1674) | 85 (0.1872) | 19 (0.6129) | 20 (0.6452) | 22 (0.7097) | 22 (0.7097) |

Table 5: EmbedPVP variant prediction results across several models using ClinVar dataset for the novel genes or diseases

| | | Novel Diseases and Known Genes | | | | Known Genes and Diseases | | | |
|---|---|---|---|---|---|---|---|---|---|
| | | **H@1** | **H@10** | **H@30** | **H@50** | **H@1** | **H@10** | **H@30** | **H@50** |
| **Genotype-based prediction tools** | **CADD** | 27 (0.2432) | 57 (0.5135) | 69 (0.6216) | 75 (0.6757) | 135 (0.2789) | 273 (0.5640) | 305 (0.6302) | 328 (0.6777) |
| | **MCAP** | 2 (0.0180) | 13 (0.1171) | 23 (0.2072) | 28 (0.2523) | 6 (0.0124) | 74 (0.1529) | 95 (0.1963) | 107 (0.2211) |
| | **SIFT** | 17 (0.1532) | 17 (0.1532) | 17 (0.1532) | 17 (0.1532) | 72 (0.1488) | 72 (0.1488) | 72 (0.1488) | 72 (0.1488) |
| | **PolyPhen2** | 11 (0.0991) | 11 (0.0991) | 11 (0.0991) | 16 (0.1441) | 38 (0.0785) | 38 (0.0785) | 38 (0.0785) | 61 (0.1260) |
| | **DANN** | 1 (0.0090) | 14 (0.1261) | 14 (0.1261) | 14 (0.1261) | 1 (0.0021) | 44 (0.0909) | 44 (0.0909) | 44 (0.0909) |
| | **MetaSVM** | 7 (0.0631) | 9 (0.0811) | 18 (0.1622) | 22 (0.1982) | 9 (0.0186) | 34 (0.0702) | 72 (0.1488) | 89 (0.1839) |
| **Phenotype-based prediction tools** | **PHIVE** | 33 (0.2973) | 38 (0.3423) | 41 (0.3694) | 42 (0.3784) | 105 (0.2169) | 183 (0.3781) | 194 (0.4008) | 196 (0.4050) |
| | **DeepPVP** | 37 (0.3333) | 73 (0.6577) | 75 (0.6757) | 76 (0.6847) | 299 (0.6178) | 371 (0.7665) | 385 (0.7955) | 389 (0.8037) |
| | **Phenix** | 4 (0.0360) | 51 (0.4595) | 80 (0.7207) | 81 (0.7297) | 93 (0.1921) | 408 (0.8430) | 427 (0.8822) | 432 (0.8926) |
| | **hiPHIVE** | **56 (0.5045)** | 71 (0.6396) | 78 (0.7027) | 80 (0.7207) | 333 (0.6880) | 389 (0.8037) | 417 (0.8616) | 431 (0.8905) |
| **EmbedPVP (ConvE)** | **GO** | 9 (0.0811) | 32 (0.2883) | 50 (0.4505) | 54 (0.4865) | 62 (0.1281) | 148 (0.3058) | 201 (0.4153) | 220 (0.4545) |
| | **HP** | 27 (0.2432) | 57 (0.5135) | 69 (0.6216) | 75 (0.6757) | 128 (0.2645) | 270 (0.5579) | 303 (0.6260) | 328 (0.6777) |
| | **MP** | 22 (0.1982) | 53 (0.4775) | 66 (0.5946) | 70 (0.6306) | 126 (0.2603) | 255 (0.5269) | 298 (0.6157) | 330 (0.6818) |
| | **UBERON** | 23 (0.2072) | 57 (0.5135) | 69 (0.6216) | 75 (0.6757) | 116 (0.2397) | 273 (0.5640) | 305 (0.6302) | 328 (0.6777) |
| | **Union** | 19 (0.1712) | 45 (0.4054) | 65 (0.5856) | 73 (0.6577) | 112 (0.2314) | 226 (0.4669) | 291 (0.6012) | 326 (0.6736) |
| **EmbedPVP (DistMult)** | **GO** | 31 (0.2793) | 61 (0.5495) | 82 (0.7387) | 87 (0.7838) | 242 (0.5000) | 326 (0.6736) | 383 (0.7913) | 411 (0.8492) |
| | **HP** | 26 (0.2342) | 70 (0.6306) | 88 (0.7928) | 93 (0.8378) | 276 (0.5702) | 427 (0.8822) | 443 (0.9153) | 449 (0.9277) |
| | **MP** | 41 (0.3694) | 58 (0.5225) | 77 (0.6937) | 85 (0.7658) | 243 (0.5021) | 309 (0.6384) | 379 (0.7831) | 411 (0.8492) |
| | **UBERON** | 21 (0.1892) | 38 (0.3423) | 54 (0.4865) | 67 (0.6036) | 154 (0.3182) | 286 (0.5909) | 343 (0.7087) | 377 (0.7789) |
| | **Union** | 30 (0.2703) | 66 (0.5946) | 81 (0.7297) | 89 (0.8018) | 191 (0.3946) | 358 (0.7397) | 419 (0.8657) | 433 (0.8946) |
| **EmbedPVP (ELEmbedding)** | **GO** | 26 (0.2342) | 50 (0.4505) | 68 (0.6126) | 75 (0.6757) | 120 (0.2479) | 249 (0.5145) | 304 (0.6281) | 330 (0.6818) |
| | **HP** | 26 (0.2342) | 62 (0.5586) | 73 (0.6577) | 79 (0.7117) | 125 (0.2583) | 293 (0.6054) | 320 (0.6612) | 361 (0.7459) |
| | **MP** | 23 (0.2072) | 58 (0.5225) | 69 (0.6216) | 77 (0.6937) | 116 (0.2397) | 281 (0.5806) | 304 (0.6281) | 343 (0.7087) |
| | **UBERON** | 24 (0.2162) | 54 (0.4865) | 69 (0.6216) | 76 (0.6847) | 117 (0.2417) | 252 (0.5207) | 303 (0.6260) | 329 (0.6798) |
| | **Union** | 23 (0.2072) | 50 (0.4505) | 68 (0.6126) | 75 (0.6757) | 118 (0.2438) | 246 (0.5083) | 304 (0.6281) | 328 (0.6777) |
| **EmbedPVP (Elboxembeddings)** | **GO** | 22 (0.1982) | 51 (0.4595) | 67 (0.6036) | 75 (0.6757) | 116 (0.2397) | 253 (0.5227) | 305 (0.6302) | 331 (0.6839) |
| | **HP** | 22 (0.1982) | 62 (0.5586) | 73 (0.6577) | 79 (0.7117) | 120 (0.2479) | 283 (0.5847) | 319 (0.6591) | 353 (0.7293) |
| | **MP** | 26 (0.2342) | 57 (0.5135) | 67 (0.6036) | 77 (0.6937) | 119 (0.2459) | 273 (0.5640) | 305 (0.6302) | 342 (0.7066) |
| | **UBERON** | 24 (0.2162) | 49 (0.4414) | 67 (0.6036) | 74 (0.6667) | 120 (0.2479) | 243 (0.5021) | 304 (0.6281) | 328 (0.6777) |
| | **Union** | 21 (0.1892) | 48 (0.4324) | 67 (0.6036) | 74 (0.6667) | 118 (0.2438) | 239 (0.4938) | 304 (0.6281) | 326 (0.6736) |
| **EmbedPVP (TransD)** | **GO** | 49 (0.4414) | 68 (0.6126) | 83 (0.7477) | 86 (0.7748) | 346 (0.7149) | 421 (0.8698) | 455 (0.9401) | 458 (0.9463) |
| | **HP** | 47 (0.4234) | **84 (0.7568)** | **91 (0.8198)** | **95 (0.8559)** | **436 (0.9008)** | **472 (0.9752)** | **476 (0.9835)** | **476 (0.9835)** |
| | **MP** | 52 (0.4685) | 78 (0.7027) | 87 (0.7838) | 90 (0.8108) | 409 (0.8450) | 455 (0.9401) | 468 (0.9669) | 470 (0.9711) |
| | **UBERON** | 51 (0.4595) | 71 (0.6396) | 86 (0.7748) | 91 (0.8198) | 397 (0.8202) | 450 (0.9298) | 467 (0.9649) | 471 (0.9731) |
| | **Union** | 44 (0.3964) | 83 (0.7477) | **91 (0.8198)** | 93 (0.8378) | 415 (0.8574) | 457 (0.9442) | 464 (0.9587) | 468 (0.9669) |
| **EmbedPVP (TransE)** | **GO** | 27 (0.2432) | 56 (0.5045) | 69 (0.6216) | 75 (0.6757) | 135 (0.2789) | 272 (0.5620) | 305 (0.6302) | 328 (0.6777) |
| | **HP** | 31 (0.2793) | 58 (0.5225) | 70 (0.6306) | 78 (0.7027) | 248 (0.5124) | 346 (0.7149) | 407 (0.8409) | 434 (0.8967) |
| | **MP** | 27 (0.2432) | 56 (0.5045) | 69 (0.6216) | 75 (0.6757) | 135 (0.2789) | 273 (0.5640) | 305 (0.6302) | 328 (0.6777) |
| | **UBERON** | 26 (0.2342) | 56 (0.5045) | 68 (0.6126) | 75 (0.6757) | 135 (0.2789) | 272 (0.5620) | 305 (0.6302) | 328 (0.6777) |
| | **Union** | 26 (0.2342) | 57 (0.5135) | 69 (0.6216) | 75 (0.6757) | 135 (0.2789) | 273 (0.5640) | 306 (0.6322) | 329 (0.6798) |
| **EmbedPVP (TransR)** | **GO** | 20 (0.1802) | 45 (0.4054) | 65 (0.5856) | 73 (0.6577) | 111 (0.2293) | 241 (0.4979) | 305 (0.6302) | 330 (0.6818) |
| | **HP** | 27 (0.2432) | 57 (0.5135) | 74 (0.6667) | 81 (0.7297) | 244 (0.5041) | 337 (0.6963) | 390 (0.8058) | 424 (0.8760) |
| | **MP** | 23 (0.2072) | 50 (0.4505) | 66 (0.5946) | 76 (0.6847) | 117 (0.2417) | 262 (0.5413) | 311 (0.6426) | 334 (0.6901) |
| | **UBERON** | 24 (0.2162) | 51 (0.4595) | 67 (0.6036) | 75 (0.6757) | 128 (0.2645) | 257 (0.5310) | 310 (0.6405) | 339 (0.7004) |
| | **Union** | 25 (0.2252) | 55 (0.4955) | 69 (0.6216) | 78 (0.7027) | 159 (0.3285) | 289 (0.5971) | 320 (0.6612) | 370 (0.7645) |
| **EmbedPVP (DL2vec)** | **GO** | 18 (0.1622) | 38 (0.3423) | 61 (0.5495) | 68 (0.6126) | 338 (0.6983) | 429 (0.8864) | 445 (0.9194) | 451 (0.9318) |
| | **HP** | 36 (0.3243) | 55 (0.4955) | 62 (0.5586) | 67 (0.6036) | 434 (0.8967) | 466 (0.9628) | 469 (0.9690) | 470 (0.9711) |
| | **MP** | 17 (0.1532) | 39 (0.3514) | 59 (0.5315) | 67 (0.6036) | 370 (0.7645) | 458 (0.9463) | 468 (0.9669) | 470 (0.9711) |
| | **UBERON** | 25 (0.2252) | 51 (0.4595) | 66 (0.5946) | 71 (0.6396) | 345 (0.7128) | 445 (0.9194) | 459 (0.9483) | 466 (0.9628) |
| | **Union** | 31 (0.2793) | 55 (0.4955) | 65 (0.5856) | 72 (0.6486) | 411 (0.8492) | 459 (0.9483) | 467 (0.9649) | 470 (0.9711) |
| **EmbedPVP (OWL2vec*)** | **GO** | 33 (0.2973) | 62 (0.5586) | 70 (0.6306) | 73 (0.6577) | 359 (0.7417) | 437 (0.9029) | 455 (0.9401) | 462 (0.9545) |
| | **HP** | 40 (0.3604) | 54 (0.4865) | 67 (0.6036) | 78 (0.7027) | 434 (0.8967) | 463 (0.9566) | 467 (0.9649) | 469 (0.9690) |
| | **MP** | 20 (0.1802) | 47 (0.4234) | 68 (0.6126) | 74 (0.6667) | 397 (0.8202) | 457 (0.9442) | 469 (0.9690) | 473 (0.9773) |
| | **UBERON** | 23 (0.2072) | 52 (0.4685) | 63 (0.5676) | 71 (0.6396) | 356 (0.7355) | 441 (0.9112) | 461 (0.9525) | 470 (0.9711) |
| | **Union** | 32 (0.2883) | 63 (0.5676) | 80 (0.7207) | 86 (0.7748) | 424 (0.8760) | 461 (0.9525) | 473 (0.9773) | **476 (0.9835)** |

Table 6: EmbedPVP variant prediction results across several models using ClinVar dataset for either known genes and/or diseases during training.

| | | Exonic | | | | Non-Exonic | | | |
|---|---|---|---|---|---|---|---|---|---|
| | | **H@1** | **H@10** | **H@30** | **H@50** | **H@1** | **H@10** | **H@30** | **H@50** |
| **Genotype-based prediction tools** | **CADD** | 239 (0.2701) | 364 (0.4113) | 448 (0.5062) | 533 (0.6023) | 0 (0.0) | 141 (0.7157) | 152 (0.7716) | 155 (0.7868) |
| | **MCAP** | 11 (0.0124) | 124 (0.1401) | 200 (0.2260) | 243 (0.2746) | 0 (0.0) | 1 (0.0051) | 1 (0.0051) | 1 (0.0051) |
| | **SIFT** | 198 (0.2237) | 198 (0.2237) | 198 (0.2237) | 198 (0.2237) | 0 (0.0) | 0 (0.0) | 0 (0.0) | 0 (0.0) |
| | **PolyPhen2** | 112 (0.1266) | 112 (0.1266) | 112 (0.1266) | 182 (0.2056) | 0 (0.0) | 0 (0.0) | 0 (0.0) | 0 (0.0) |
| | **DANN** | 7 (0.0079) | 172 (0.1944) | 172 (0.1944) | 172 (0.1944) | 0 (0.0) | 0 (0.0) | 0 (0.0) | 0 (0.0) |
| | **MetaSVM** | 20 (0.0226) | 74 (0.0836) | 139 (0.1571) | 177 (0.2000) | 0 (0.0) | 0 (0.0) | 0 (0.0) | 0 (0.0) |
| **Phenotype-based prediction tools** | **PHIVE** | 129 (0.1458) | 208 (0.2350) | 218 (0.2463) | 220 (0.2486) | 46 (0.2335) | 76 (0.3858) | 80 (0.4061) | 81 (0.4112) |
| | **DeepPVP** | 328 (0.3706) | 422 (0.4768) | 444 (0.5017) | 453 (0.5119) | 68 (0.3452) | 110 (0.5584) | 116 (0.5888) | 119 (0.6041) |
| | **Phenix** | 98 (0.1107) | 441 (0.4983) | 479 (0.5412) | 490 (0.5537) | 20 (0.1015) | 126 (0.6396) | 141 (0.7157) | 143 (0.7259) |
| | **hiPHIVE** | 355 (0.4011) | 447 (0.5051) | 499 (0.5638) | 519 (0.5864) | **132 (0.6701)** | 139 (0.7056) | 142 (0.7208) | 143 (0.7259) |
| **EmbedPVP (ConvE)** | **GO** | 100 (0.1130) | 221 (0.2497) | 309 (0.3492) | 347 (0.3921) | 2 (0.0102) | 51 (0.2589) | 76 (0.3858) | 88 (0.4467) |
| | **HP** | 220 (0.2486) | 364 (0.4113) | 447 (0.5051) | 533 (0.6023) | 0 (0.0) | 137 (0.6954) | 150 (0.7614) | 153 (0.7766) |
| | **MP** | 203 (0.2294) | 349 (0.3944) | 438 (0.4949) | 516 (0.5831) | 5 (0.0254) | 111 (0.5635) | 141 (0.7157) | 146 (0.7411) |
| | **UBERON** | 205 (0.2316) | 364 (0.4113) | 447 (0.5051) | 533 (0.6023) | 0 (0.0) | 140 (0.7107) | 152 (0.7716) | 155 (0.7868) |
| | **Union** | 186 (0.2102) | 320 (0.3616) | 426 (0.4814) | 514 (0.5808) | 2 (0.0102) | 81 (0.4112) | 138 (0.7005) | 148 (0.7513) |
| **EmbedPVP (DistMult)** | **GO** | 255 (0.2881) | 371 (0.4192) | 491 (0.5548) | 549 (0.6203) | 59 (0.2995) | 109 (0.5533) | 137 (0.6954) | 145 (0.7360) |
| | **HP** | 262 (0.2960) | 451 (0.5096) | 481 (0.5435) | 489 (0.5525) | 83 (0.4213) | 120 (0.6091) | 130 (0.6599) | 134 (0.6802) |
| | **MP** | 287 (0.3243) | 374 (0.4226) | 522 (0.5898) | 594 (0.6712) | 63 (0.3198) | 115 (0.5838) | 141 (0.7157) | 150 (0.7614) |
| | **UBERON** | 201 (0.2271) | 346 (0.3910) | 468 (0.5288) | 545 (0.6158) | 13 (0.0660) | 115 (0.5838) | 140 (0.7107) | 144 (0.7310) |
| | **Union** | 208 (0.2350) | 383 (0.4328) | 490 (0.5537) | 528 (0.5966) | 38 (0.1929) | 115 (0.5838) | 126 (0.6396) | 133 (0.6751) |
| **EmbedPVP (ELEmbedding)** | **GO** | 215 (0.2429) | 350 (0.3955) | 448 (0.5062) | 540 (0.6102) | 0 (0.0) | 104 (0.5279) | 152 (0.7716) | 155 (0.7868) |
| | **HP** | 189 (0.2136) | 378 (0.4271) | 461 (0.5209) | 538 (0.6079) | 0 (0.0) | 137 (0.6954) | 155 (0.7868) | 156 (0.7919) |
| | **MP** | 209 (0.2362) | 378 (0.4271) | 445 (0.5028) | 551 (0.6226) | 0 (0.0) | 140 (0.7107) | 152 (0.7716) | 155 (0.7868) |
| | **UBERON** | 205 (0.2316) | 355 (0.4011) | 446 (0.5040) | 537 (0.6068) | 0 (0.0) | 111 (0.5635) | 152 (0.7716) | 154 (0.7817) |
| | **Union** | 206 (0.2328) | 351 (0.3966) | 443 (0.5006) | 536 (0.6056) | 0 (0.0) | 103 (0.5228) | 152 (0.7716) | 155 (0.7868) |
| **EmbedPVP (Elboxembeddings)** | **GO** | 202 (0.2282) | 353 (0.3989) | 447 (0.5051) | 538 (0.6079) | 0 (0.0) | 111 (0.5635) | 152 (0.7716) | 155 (0.7868) |
| | **HP** | 180 (0.2034) | 374 (0.4226) | 459 (0.5186) | 531 (0.6000) | 0 (0.0) | 131 (0.6650) | 155 (0.7868) | 156 (0.7919) |
| | **MP** | 208 (0.2350) | 369 (0.4169) | 450 (0.5085) | 548 (0.6192) | 0 (0.0) | 131 (0.6650) | 152 (0.7716) | 155 (0.7868) |
| | **UBERON** | 213 (0.2407) | 349 (0.3944) | 438 (0.4949) | 531 (0.6000) | 0 (0.0) | 93 (0.4721) | 152 (0.7716) | 154 (0.7817) |
| | **Union** | 204 (0.2305) | 348 (0.3932) | 447 (0.5051) | 529 (0.5977) | 0 (0.0) | 92 (0.4670) | 152 (0.7716) | 155 (0.7868) |
| **EmbedPVP (TransD)** | **GO** | 366 (0.4136) | 506 (0.5718) | **632 (0.7141)** | **680 (0.7684)** | 97 (0.4924) | 140 (0.7107) | 166 (0.8426) | 166 (0.8426) |
| | **HP** | **438 (0.4949)** | 494 (0.5582) | 503 (0.5684) | 515 (0.5819) | 116 (0.5888) | 139 (0.7056) | 144 (0.7310) | 145 (0.7360) |
| | **MP** | 417 (0.4712) | 540 (0.6102) | 619 (0.6994) | 653 (0.7379) | 109 (0.5533) | 141 (0.7157) | 158 (0.8020) | 162 (0.8223) |
| | **UBERON** | 379 (0.4282) | 503 (0.5684) | 590 (0.6667) | 640 (0.7232) | 106 (0.5381) | 130 (0.6599) | 158 (0.8020) | 168 (0.8528) |
| | **Union** | 426 (0.4814) | **548 (0.6192)** | 604 (0.6825) | 637 (0.7198) | 111 (0.5635) | 143 (0.7259) | **164 (0.8325)** | **171 (0.8680)** |
| **EmbedPVP (TransE)** | **GO** | 232 (0.2621) | 361 (0.4079) | 448 (0.5062) | 533 (0.6023) | 0 (0.0) | 128 (0.6497) | 152 (0.7716) | 155 (0.7868) |
| | **HP** | 308 (0.3480) | 439 (0.4960) | 567 (0.6407) | 653 (0.7379) | 66 (0.3350) | 154 (0.7817) | 162 (0.8223) | 167 (0.8477) |
| | **MP** | 232 (0.2621) | 358 (0.4045) | 448 (0.5062) | 533 (0.6023) | 0 (0.0) | 128 (0.6497) | 152 (0.7716) | 155 (0.7868) |
| | **UBERON** | 226 (0.2554) | 359 (0.4056) | 444 (0.5017) | 533 (0.6023) | 0 (0.0) | 125 (0.6345) | 152 (0.7716) | 155 (0.7868) |
| | **Union** | 231 (0.2610) | 361 (0.4079) | 449 (0.5073) | 534 (0.6034) | 0 (0.0) | 135 (0.6853) | 152 (0.7716) | 155 (0.7868) |
| **EmbedPVP (TransR)** | **GO** | 204 (0.2305) | 356 (0.4023) | 451 (0.5096) | 537 (0.6068) | 0 (0.0) | 97 (0.4924) | 152 (0.7716) | 155 (0.7868) |
| | **HP** | 313 (0.3537) | 432 (0.4881) | 549 (0.6203) | 648 (0.7322) | 56 (0.2843) | **146 (0.7411)** | 160 (0.8122) | 164 (0.8325) |
| | **MP** | 200 (0.2260) | 362 (0.4090) | 455 (0.5141) | 545 (0.6158) | 0 (0.0) | 112 (0.5685) | 152 (0.7716) | 155 (0.7868) |
| | **UBERON** | 219 (0.2475) | 362 (0.4090) | 452 (0.5107) | 546 (0.6169) | 2 (0.0102) | 106 (0.5381) | 152 (0.7716) | 155 (0.7868) |
| | **Union** | 251 (0.2836) | 376 (0.4249) | 469 (0.5299) | 586 (0.6621) | 13 (0.0660) | 137 (0.6954) | 154 (0.7817) | 158 (0.8020) |
| **EmbedPVP (DL2vec)** | **GO** | 327 (0.3695) | 432 (0.4881) | 486 (0.5492) | 515 (0.5819) | 87 (0.4416) | 122 (0.6193) | 137 (0.6954) | 143 (0.7259) |
| | **HP** | 428 (0.4836) | 467 (0.5277) | 474 (0.5356) | 479 (0.5412) | 114 (0.5787) | 130 (0.6599) | 133 (0.6751) | 134 (0.6802) |
| | **MP** | 338 (0.3819) | 451 (0.5096) | 499 (0.5638) | 519 (0.5864) | 103 (0.5228) | 125 (0.6345) | 133 (0.6751) | 137 (0.6954) |
| | **UBERON** | 332 (0.3751) | 450 (0.5085) | 501 (0.5661) | 530 (0.5989) | 81 (0.4112) | 124 (0.6294) | 141 (0.7157) | 145 (0.7360) |
| | **Union** | 403 (0.4554) | 477 (0.5390) | 503 (0.5684) | 525 (0.5932) | 108 (0.5482) | 128 (0.6497) | 135 (0.6853) | 144 (0.7310) |
| **EmbedPVP (OWL2vec*)** | **GO** | 355 (0.4011) | 477 (0.5390) | 540 (0.6102) | 558 (0.6305) | 101 (0.5127) | 132 (0.6701) | 140 (0.7107) | 145 (0.7360) |
| | **HP** | 432 (0.4881) | 467 (0.5277) | 479 (0.5412) | 486 (0.5492) | 115 (0.5838) | 128 (0.6497) | 133 (0.6751) | 139 (0.7056) |
| | **MP** | 373 (0.4215) | 466 (0.5266) | 516 (0.5831) | 542 (0.6124) | 110 (0.5584) | 134 (0.6802) | 144 (0.7310) | 149 (0.7563) |
| | **UBERON** | 335 (0.3785) | 450 (0.5085) | 491 (0.5548) | 514 (0.5808) | 91 (0.4619) | 126 (0.6396) | 133 (0.6751) | 141 (0.7157) |
| | **Union** | 411 (0.4644) | 479 (0.5412) | 518 (0.5853) | 533 (0.6023) | 116 (0.5888) | 131 (0.6650) | 143 (0.7259) | 146 (0.7411) |

Table 7: EmbedPVP variant prediction results across several models using ClinVar dataset for the exonic and non-exonic variants

##### 2.1.4 Evaluations for the variants in genes with no phenotype annotations

Table 8.

| | | H@1 | H@10 | H@30 | H@50 |
|---|---|---|---|---|---|
| Genotype-based prediction tools | **CADD** | 71 (0.1788) | 145 (0.3652) | 189 (0.4761) | 239 (0.6020) |
| | **MCAP** | 1 (0.0025) | 32 (0.0806) | 66 (0.1662) | 86 (0.2166) |
| | **SIFT** | **95 (0.2393)** | 95 (0.2393) | 95 (0.2393) | 95 (0.2393) |
| | **PolyPhen2** | 45 (0.1134) | 45 (0.1134) | 45 (0.1134) | 82 (0.2065) |
| | **DANN** | 5 (0.0126) | 96 (0.2418) | 96 (0.2418) | 96 (0.2418) |
| | **MetaSVM** | 4 (0.0101) | 25 (0.0630) | 39 (0.0982) | 53 (0.1335) |
| Phenotype-based prediction tools | **PHIVE** | 24 (0.0605) | 41 (0.1033) | 42 (0.1058) | 42 (0.1058) |
| | **DeepPVP** | 36 (0.0907) | 53 (0.1335) | 61 (0.1537) | 68 (0.1713) |
| | **Phenix** | 8 (0.0202) | 63 (0.1587) | 68 (0.1713) | 72 (0.1814) |
| | **hiPHIVE** | 63 (0.1587) | 89 (0.2242) | 103 (0.2594) | 105 (0.2645) |
| EmbedPVP (ConvE) | **GO** | 28 (0.0705) | 83 (0.2091) | 120 (0.3023) | 141 (0.3552) |
| | **HP** | 61 (0.1537) | 145 (0.3652) | 189 (0.4761) | 238 (0.5995) |
| | **MP** | 56 (0.1411) | 132 (0.3325) | 183 (0.4610) | 225 (0.5668) |
| | **UBERON** | 60 (0.1511) | 145 (0.3652) | 188 (0.4736) | 239 (0.6020) |
| | **Union** | 50 (0.1259) | 113 (0.2846) | 173 (0.4358) | 221 (0.5567) |
| EmbedPVP (DistMult) | **GO** | 43 (0.1083) | 84 (0.2116) | 140 (0.3526) | 164 (0.4131) |
| | **HP** | 2 (0.0050) | 8 (0.0202) | 11 (0.0277) | 13 (0.0327) |
| | **MP** | 57 (0.1436) | 107 (0.2695) | 177 (0.4458) | 214 (0.5390) |
| | **UBERON** | 36 (0.0907) | 116 (0.2922) | 180 (0.4534) | 208 (0.5239) |
| | **Union** | 9 (0.0227) | 36 (0.0907) | 65 (0.1637) | 84 (0.2116) |
| EmbedPVP (ELEmbedding) | **GO** | 64 (0.1612) | 133 (0.3350) | 191 (0.4811) | 242 (0.6096) |
| | **HP** | 34 (0.0856) | 128 (0.3224) | 178 (0.4484) | 203 (0.5113) |
| | **MP** | 64 (0.1612) | **153 (0.3854)** | 187 (0.4710) | 236 (0.5945) |
| | **UBERON** | 58 (0.1461) | 135 (0.3401) | 189 (0.4761) | 238 (0.5995) |
| | **Union** | 59 (0.1486) | 133 (0.3350) | 186 (0.4685) | 241 (0.6071) |
| EmbedPVP (Elboxembeddings) | **GO** | 60 (0.1511) | 135 (0.3401) | 189 (0.4761) | 239 (0.6020) |
| | **HP** | 33 (0.0831) | 128 (0.3224) | 178 (0.4484) | 205 (0.5164) |
| | **MP** | 58 (0.1461) | 144 (0.3627) | 192 (0.4836) | 236 (0.5945) |
| | **UBERON** | 64 (0.1612) | 129 (0.3249) | 182 (0.4584) | 237 (0.5970) |
| | **Union** | 59 (0.1486) | 130 (0.3275) | 191 (0.4811) | 236 (0.5945) |
| EmbedPVP (TransD) | **GO** | 64 (0.1612) | 138 (0.3476) | 220 (0.5542) | 249 (0.6272) |
| | **HP** | 19 (0.0479) | 27 (0.0680) | 29 (0.0730) | 34 (0.0856) |
| | **MP** | 61 (0.1537) | 133 (0.3350) | **196 (0.4937)** | 218 (0.5491) |
| | **UBERON** | 46 (0.1159) | 107 (0.2695) | 174 (0.4383) | 211 (0.5315) |
| | **Union** | 20 (0.0504) | 87 (0.2191) | 144 (0.3627) | 175 (0.4408) |
| EmbedPVP (TransE) | **GO** | 64 (0.1612) | 136 (0.3426) | 189 (0.4761) | 239 (0.6020) |
| | **HP** | 65 (0.1637) | 143 (0.3602) | 189 (0.4761) | 239 (0.6020) |
| | **MP** | 66 (0.1662) | 133 (0.3350) | 189 (0.4761) | 239 (0.6020) |
| | **UBERON** | 60 (0.1511) | 132 (0.3325) | 186 (0.4685) | 239 (0.6020) |
| | **Union** | 63 (0.1587) | 137 (0.3451) | 189 (0.4761) | 239 (0.6020) |
| EmbedPVP (TransR) | **GO** | 67 (0.1688) | 145 (0.3652) | 193 (0.4861) | **243 (0.6121)** |
| | **HP** | 63 (0.1587) | 138 (0.3476) | 185 (0.4660) | 239 (0.6020) |
| | **MP** | 52 (0.1310) | 136 (0.3426) | 192 (0.4836) | 240 (0.6045) |
| | **UBERON** | 63 (0.1587) | 137 (0.3451) | 190 (0.4786) | 238 (0.5995) |
| | **Union** | 62 (0.1562) | 135 (0.3401) | 190 (0.4786) | 241 (0.6071) |
| EmbedPVP (DL2vec) | **GO** | 19 (0.0479) | 41 (0.1033) | 67 (0.1688) | 88 (0.2217) |
| | **HP** | 18 (0.0453) | 21 (0.0529) | 23 (0.0579) | 23 (0.0579) |
| | **MP** | 18 (0.0453) | 36 (0.0907) | 55 (0.1385) | 67 (0.1688) |
| | **UBERON** | 23 (0.0579) | 42 (0.1058) | 72 (0.1814) | 88 (0.2217) |
| | **Union** | 17 (0.0428) | 38 (0.0957) | 54 (0.1360) | 73 (0.1839) |
| EmbedPVP (OWL2vec*) | **GO** | 26 (0.0655) | 61 (0.1537) | 104 (0.2620) | 117 (0.2947) |
| | **HP** | 19 (0.0479) | 23 (0.0579) | 24 (0.0605) | 24 (0.0605) |
| | **MP** | 21 (0.0529) | 47 (0.1184) | 70 (0.1763) | 89 (0.2242) |
| | **UBERON** | 20 (0.0504) | 32 (0.0806) | 49 (0.1234) | 60 (0.1511) |
| | **Union** | 15 (0.0378) | 31 (0.0781) | 50 (0.1259) | 60 (0.1511) |

Table 8: EmbedPVP evaluations for the variants in genes with no phenotype annotations

##### 2.1.5 Evaluations for the variants in intergenic and overlapping genes

Table 9.

| | | H@1 | H@10 | H@30 | H@50 |
|---|---|---|---|---|---|
| Genotype-based prediction tools | CADD | 30 (0.2778) | 53 (0.4907) | 59 (0.5463) | 66 (0.6111) |
| | MCAP | 3 (0.0278) | 7 (0.0648) | 14 (0.1296) | 18 (0.1667) |
| | SIFT | 14 (0.1296) | 14 (0.1296) | 14 (0.1296) | 14 (0.1296) |
| | PolyPhen2 | 13 (0.1204) | 13 (0.1204) | 13 (0.1204) | 16 (0.1481) |
| | DANN | 0 (0.0) | 12 (0.1111) | 12 (0.1111) | 12 (0.1111) |
| | MetaSVM | 0 (0.0) | 4 (0.0370) | 14 (0.1296) | 16 (0.1481) |
| Phenotype-based prediction tools | PHIVE | 18 (0.1667) | 40 (0.3704) | 42 (0.3889) | 42 (0.3889) |
| | DeepPVP | 42 (0.3889) | 60 (0.5556) | 64 (0.5926) | 65 (0.6019) |
| | Phenix | 17 (0.1574) | 72 (0.6667) | 75 (0.6944) | 78 (0.7222) |
| | hiPHIVE | 59 (0.5463) | 63 (0.5833) | 66 (0.6111) | 68 (0.6296) |
| EmbedPVP (ConvE) | GO | 17 (0.1574) | 36 (0.3333) | 47 (0.4352) | 51 (0.4722) |
| | HP | 29 (0.2685) | 52 (0.4815) | 59 (0.5463) | 66 (0.6111) |
| | MP | 30 (0.2778) | 54 (0.5) | 57 (0.5278) | 62 (0.5741) |
| | UBERON | 28 (0.2593) | 53 (0.4907) | 59 (0.5463) | 66 (0.6111) |
| | Union | 30 (0.2778) | 44 (0.4074) | 57 (0.5278) | 64 (0.5926) |
| EmbedPVP (DistMult) | GO | 41 (0.3796) | 55 (0.5093) | 68 (0.6296) | 72 (0.6667) |
| | HP | 45 (0.4167) | 71 (0.6574) | 74 (0.6852) | 77 (0.7130) |
| | MP | 42 (0.3889) | 55 (0.5093) | 65 (0.6019) | 71 (0.6574) |
| | UBERON | 27 (0.25) | 48 (0.4444) | 59 (0.5463) | 71 (0.6574) |
| | Union | 40 (0.3704) | 58 (0.5370) | 71 (0.6574) | 76 (0.7037) |
| EmbedPVP (ELEmbedding) | GO | 28 (0.2593) | 47 (0.4352) | 59 (0.5463) | 67 (0.6204) |
| | HP | 28 (0.2593) | 57 (0.5278) | 65 (0.6019) | 69 (0.6389) |
| | MP | 28 (0.2593) | 54 (0.5000) | 59 (0.5463) | 70 (0.6481) |
| | UBERON | 28 (0.2593) | 47 (0.4352) | 59 (0.5463) | 69 (0.6389) |
| | Union | 29 (0.2685) | 48 (0.4444) | 59 (0.5463) | 67 (0.6204) |
| EmbedPVP (Elboxembeddings) | GO | 27 (0.25) | 51 (0.4722) | 59 (0.5463) | 68 (0.6296) |
| | HP | 27 (0.25) | 55 (0.5093) | 64 (0.5926) | 68 (0.6296) |
| | MP | 29 (0.2685) | 53 (0.4907) | 59 (0.5463) | 68 (0.6296) |
| | UBERON | 28 (0.2593) | 49 (0.4537) | 59 (0.5463) | 65 (0.6019) |
| | Union | 29 (0.2685) | 47 (0.4352) | 59 (0.5463) | 67 (0.6204) |
| EmbedPVP (TransD) | GO | 61 (0.5648) | 73 (0.6759) | 87 (0.8056) | 89 (0.8241) |
| | HP | 65 (0.6019) | 76 (0.7037) | 78 (0.7222) | 78 (0.7222) |
| | MP | 63 (0.5833) | 71 (0.6574) | 79 (0.7315) | 83 (0.7685) |
| | UBERON | 61 (0.5648) | 69 (0.6389) | 73 (0.6759) | 78 (0.7222) |
| | Union | 65 (0.6019) | **79 (0.7315)** | **84 (0.7778)** | **87 (0.8056)** |
| EmbedPVP (TransE) | GO | 30 (0.2778) | 52 (0.4815) | 59 (0.5463) | 66 (0.6111) |
| | HP | 42 (0.3889) | 66 (0.6111) | 73 (0.6759) | 78 (0.7222) |
| | MP | 29 (0.2685) | 52 (0.4815) | 59 (0.5463) | 66 (0.6111) |
| | UBERON | 29 (0.2685) | 52 (0.4815) | 59 (0.5463) | 66 (0.6111) |
| | Union | 30 (0.2778) | 53 (0.4907) | 59 (0.5463) | 66 (0.6111) |
| EmbedPVP (TransR) | GO | 28 (0.2593) | 47 (0.4352) | 60 (0.5556) | 65 (0.6019) |
| | HP | 42 (0.3889) | 62 (0.5741) | 70 (0.6481) | 77 (0.7130) |
| | MP | 26 (0.2407) | 51 (0.4722) | 60 (0.5556) | 69 (0.6389) |
| | UBERON | 29 (0.2685) | 50 (0.4630) | 59 (0.5463) | 67 (0.6204) |
| | Union | 34 (0.3148) | 56 (0.5185) | 61 (0.5648) | 71 (0.6574) |
| EmbedPVP (DL2vec) | GO | 56 (0.5185) | 74 (0.6852) | 76 (0.7037) | 77 (0.7130) |
| | HP | 65 (0.6019) | 75 (0.6944) | 76 (0.7037) | 76 (0.7037) |
| | MP | 64 (0.5926) | 72 (0.6667) | 76 (0.7037) | 77 (0.7130) |
| | UBERON | 57 (0.5278) | 67 (0.6204) | 72 (0.6667) | 76 (0.7037) |
| | Union | 64 (0.5926) | 72 (0.6667) | 73 (0.6759) | 76 (0.7037) |
| EmbedPVP (OWL2vec*) | GO | 61 (0.5648) | 70 (0.6481) | 79 (0.7315) | 80 (0.7407) |
| | HP | 66 (0.6111) | 74 (0.6852) | 75 (0.6944) | 76 (0.7037) |
| | MP | 61 (0.5648) | 73 (0.6759) | 76 (0.7037) | 78 (0.7222) |
| | UBERON | 62 (0.5741) | 74 (0.6852) | 75 (0.6944) | 77 (0.7130) |
| | Union | **68 (0.6296)** | 74 (0.6852) | 80 (0.7407) | 82 (0.7593) |

Table 9: EmbedPVP evaluations for the variants within overlapping genes

#### 2.2 Evaluations for the variants in non-exonic regions

Evaluating variants in non-exonic regions is crucial as they can significantly impact gene expression and regulation. These regions contain important regulatory elements, such as enhancers and silencers, that play a crucial role in the expression of neighboring genes. By analyzing these variants, we can comprehensively understand the genetic factors that contribute to various diseases and conditions [1]. Therefore, we further extended our evaluation to include variants in non-exonic regions, specifically focusing on capturing phenotype annotations from the neighboring genes. Using the PAVS benchmark dataset, we identified a total of 296 non-exonic variants, including intronic, splicing, UTR5, UTR3, upstream, and ncRNA variants. The results are shown in Supplementary Table 2. Table 7 shows the results obtained using the ClinVar dataset with 197 non-exonic variants. The results highlight that our EmbedPVP models continue to outperform other state-of-the-art methods, even when considering variants in these non-exonic regions.

#### 2.3 Evaluations for variants in intergenic and overlapping genes

To further assess the performance of EmbedPVP in capturing variants located in overlapping or intergenic regions, we used the maximum similarity score among the genes surrounding the variants. To evaluate the effectiveness of our approach, we collected a new set of variants (108 variants) from ClinVar, in which the variants are within the intergenic region or overlapping genes. The results presented in Supplementary Table 9 demonstrate that our method outperforms other approaches across various metrics. Specifically, the EmbedPVP (TransD) model is considered the most effective method in capturing the genomic context and achieves better performance compared to the other methods considered in this study. Also, EmbedPVP performs better with other embedding methods, such as DL2vec and OWL2vec*, compared to the other methods.

Using this approach, in which we incorporate information for the surrounding genes, enables us to consider the genomic information surrounding the variants and thereby enhances the performance of EmbedPVP to predict other types of variants compared to other methods. By considering the genes in proximity to the variants, we ensure that the models capture the relevant genomic context necessary for accurately predicting the impact of these variants on phenotypes or diseases.

#### 2.4 Evaluations for variants in genes with no phenotype annotations

Since we utilize different types of features characterized through the use of ontologies, our method can be applied to a much larger number of genes for which the functions, sites of expression, phenotypes, or interactions with other genes are known. To evaluate our method, we focused on subsets of variants (397 variants) collected from ClinVar that corresponded to genes with no human phenotype annotations. We obtained a total of 397 variants with Gene Ontology (GO) annotations. Using the gene functions in addition to the enriched knowledge graph, we ranked the variants and assessed their performance. The results are presented in Supplementary

Table 8. Based on these results, our method, EmbedPVP, outperformed the other methods that mainly rely on existing knowledge for gene-to-disease phenotype annotations.

#### 2.5 Inductive approach

The evaluations on various benchmark datasets using the `inductive approach`. The results for PAVS, using the HP model for the clinical phenotypes and OWL2vec* embeddings, Table 10.

| | | **H@1** | **H@10** | **H@30** | **H@50** |
|---|---|---|---|---|---|
| **EmbedPVP (TransE)** | **Inductive** | 204 (0.1335) | 388 (0.2539) | 535 (0.3501) | 678 (0.4437) |
| | **Transductive** | 218 (0.1427) | 415 (0.2716) | 589 (0.3855) | 710 (0.4647) |
| **EmbedPVP (TransD)** | **Inductive** | 168 (0.1099) | 491 (0.3213) | 706 (0.4620) | 847 (0.5543) |
| | **Transductive** | 482 (0.3154) | 846 (0.5537) | 1007 (0.6590) | 1056 (0.6911) |
| **EmbedPVP (DL2vec)** | **Inductive** | 202 (0.1322) | 409 (0.2677) | 558 (0.3652) | 613 (0.4012) |
| | **Transductive** | 362 (0.2369) | 666 (0.4359) | 787 (0.5151) | 826 (0.5406) |
| **EmbedPVP (OWL2vec*)** | **Inductive** | 179 (0.1171) | 386 (0.2526) | 543 (0.3554) | 612 (0.4005) |
| | **Transductive** | 409 (0.2677) | 685 (0.4483) | 783 (0.5124) | 842 (0.5510) |

Table 10: EmbedPVP variant prediction results using for inductive vs. transductive approach for the HP model using the clinical phenotypes and selected models based on the best-performing models using the Transductive approach. The average dropped in performance using the Inductive approach for all the different models is around 12%.
